# Starvation responses impact interaction dynamics of human gut bacteria *Bacteroides thetaiotaomicron* and *Roseburia intestinalis*

**DOI:** 10.1101/2023.02.02.526806

**Authors:** Bin Liu, Daniel Rios Garza, Didier Gonze, Anna Krzynowek, Kenneth Simoens, Kristel Bernaerts, Annelies Geirnaert, Karoline Faust

**Affiliations:** Department of Microbiology, Immunology and Transplantation, Rega Institute for Medical Research, Laboratory of Molecular Bacteriology, KU Leuven, B-3000 Leuven, Belgium; Unité de Chronobiologie Théorique, Faculté des Sciences, CP 231, Université Libre de Bruxelles, Bvd du Triomphe, B-1050 Bruxelles, Belgium; Department of Chemical Engineering, Chemical and Biochemical Reactor Engineering and Safety (CREaS), KU Leuven, B-3001 Leuven, Belgium; Laboratory of Food Biotechnology, Department of Health Sciences and Technology, Institute of Food, Nutrition and Health, ETH Zürich, CH-8092 Zürich, Switzerland

## Abstract

Bacterial growth often alters the environment, which in turn can impact interspecies interactions among bacteria. Here, we used an *in vitro* batch system containing mucin beads to emulate the dynamic host environment and to study its impact on the interactions between two abundant and prevalent human gut bacteria, the primary fermenter *Bacteroides thetaiotaomicron* and the butyrate producer *Roseburia intestinalis*. By combining machine learning and flow cytometry, we found that the number of viable *B. thetaiotaomicron* cells decreases with glucose consumption due to acid production, while *R. intestinalis* survives post-glucose depletion by entering a slow growth mode. Both species attach to mucin beads, but only viable cell counts of *B. thetaiotaomicron* increase significantly. The number of viable co-culture cells varies significantly over time compared to those of monocultures. A combination of targeted metabolomics and RNA-seq showed that the slow growth mode of *R. intestinalis* represents a diauxic shift towards acetate and lactate consumption, whereas *B. thetaiotaomicron* survives glucose depletion and low pH by foraging on mucin sugars. In addition, most of the mucin monosaccharides we tested inhibited the growth of *R. intestinalis* but not *B. thetaiotaomicron*. We encoded these causal relationships in a kinetic model, which reproduced the observed dynamics. In summary, we explored how *R. intestinalis* and *B. thetaiotaomicron* respond to nutrient scarcity and how this affects their dynamics. We highlight the importance of understanding bacterial metabolic strategies to effectively modulate microbial dynamics in changing conditions.

## Introduction

The human gut is populated by hundreds of microbial species interacting with each other and with their host. Gut bacterial species differ in their distribution along the human digestive tract and favor different microhabitats, such as the lumen of the large intestine, colonic crypts and mucus layers [1, 2]. Diverse growth rates and attachment properties of gut bacteria help establish spatially structured communities. Also, the spatially heterogeneous environment influences the colonization success of different members of the gut microbiota and their competitive or cooperative relationships with one another [3–8].

An efficient way to acquire nutrients is essential for microbial persistence in the human gut. In addition to dietary fiber in the lumen, host-derived mucin glycans provide an alternative source of nutrients for members of the gut microbiota [9]. The colonic mucus layer, located at the interface between the epithelium and the gut microbiota, is key to maintain intestinal homeostasis and to enhance the resilience to perturbations of the gut microbiome [10, 11]. Colon mucus is continuously secreted by goblet cells and is organized into two distinct layers: an inner dense layer tightly attached to the epithelium, which appears to be essentially sterile when healthy; and an outer loose layer with an expanded volume, which supports a metabolically diverse community. The outer layer provides mucin polysaccharides and serves as attachment sites for the gut species that co-evolved to utilize these substrates, giving rise to complex microbial interactions [9, 10, 12, 13]. *Bacteroides* is one of the most abundant but also most variable genera across human fecal samples [14]. Its most studied species *Bacteroides thetaiotaomicron* (*B. thetaiotaomicron*) holds a large repertoire of genes involved in sensing and hydrolyzing many different polysaccharides, and *B. thetaiotaomicron* was reported to strongly express genes involved in the degradation of dietary plant glycans [15, 16]. As a nutritional generalist, *B. thetaiotaomicron* can alter its metabolic response to mucin glycans in the absence of dietary nutrients, to colonize the mucus layer and maintain its persistence in the human gut [17].

Short-chain fatty acids (SCFAs) including acetate, propionate and butyrate, are the major end products of the anaerobic fermentation of polysaccharides by the human gut microbiota. All these SCFAs exert multiple health benefits, in particular butyrate, which is used preferentially as an energy source to fuel colonic enterocytes and may trigger signals regulating mucin synthesis and secretion, as well as to maintain the integrity of epithelial barriers [18, 19]. Metabolic cross-feeding of the human gut microbiota impacts SCFA production [20]. For example, members of the most abundant genera, including *Faecalibacterium*, *Eubacterium* and *Roseburia*, are able to convert acetate and/or lactate into butyrate using the prevalent butyrate-producing pathway via butyryl-CoA:acetate CoA-transferase [21, 22]. Lactate is also an important end product of anaerobic fermentation in the gut [23]. In addition to acetate as a key intermediate that promotes cross-feeding interactions of the gut microbiota, the role of lactate as substrate for butyrate producers or for propionate producers via the acrylate pathway may explain its low levels of accumulation in the human colon of healthy adults [24, 25]. Upon exhaustion of the primary carbon sources such as carbohydrates in the gut, microbes have to hunt for another carbon source to thrive. It was reported that acetate could promote the growth of a common intestinal butyrate-producing species *Roseburia intestinalis* (*R. intestinalis*) during growth on glucose [26–28].

Several studies employing *in vitro* fermentation systems have shown mucin as a driving force in forming a microbial community that is phylogenetically and metabolically distinct from the luminal one [29, 30]. 16S rRNA gene-based results of complex communities from an *in vitro* gut model showed that *R. intestinalis* colonized mucin-covered carriers [31]. For unclear reasons, *Roseburia* but not *Bacteroides* decreased in relative abundance with mucin supplementation [32]. We thus require a mechanistic understanding of how mucin formed microhabitats impact the abundance of different gut species. Here, we built an *in vitro* batch system containing mucin beads, to systematically investigate how a spatially structured environment in the presence of mucin beads influences the growth and interactions of the model gut species *B. thetaiotaomicron* and *R. intestinalis* (Figure 1). The growth dynamics of monocultures and co-cultures in the liquid medium and attached to the beads were monitored using live/dead cell staining followed by flow cytometry in combination with machine learning techniques to quantify their proportions in co-culture. Subsequently, we characterized the effect of mucin monosaccharides on the abundance of *B. thetaiotaomicron* and *R. intestinalis*, and found that it depended on the sugar level being considered. Transcriptomic analyses of monoculture and co-culture in medium with and without mucin beads at several time points revealed different nutrient harvesting strategies employed by *B. thetaiotaomicron* and *R. intestinalis*, which resulted in their different growth modes, and also changed their interaction mechanisms depending on nutrient availability. We further developed a kinetic model that accounts for pH conditions, metabolites, and growth profiles, to mechanistically understand how gel-entrapped mucin influences the proliferation of *B. thetaiotaomicron* and *R. intestinalis* and the dynamics of their relationship. Our methodology could potentially be applied to other ecosystems where microbial growth alters environmental conditions, which in turn affect microbial interactions.

**Figure 1.**
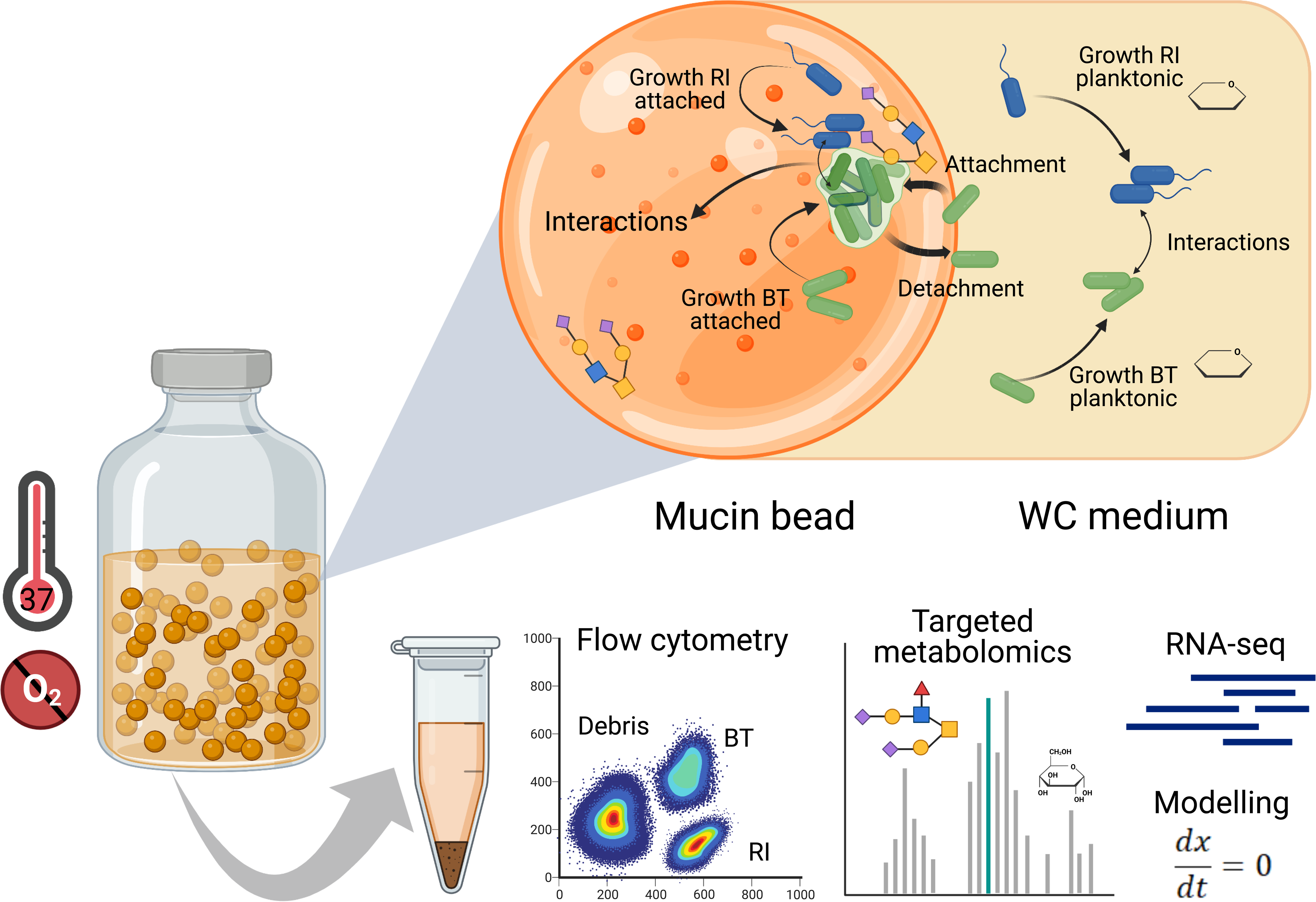
Experimental setup and design. We used a bottle system containing mucin beads in WC medium to mimic the spatially structured environment of the human gut. The growth dynamics of monoculture and co-culture in the liquid medium and attached to the beads were monitored using flow cytometry with live/dead cell staining. Live cells of *B. thetaiotaomicron* and *R. intestinalis* were differentiated from each other and from debris using a UMAP-based classification strategy (details in the “Methods” section). Fermentation products, including short-chain fatty acids and mucin-derived monomeric sugars, were measured by high-performance liquid chromatography and gas chromatography. The presence and quantity of mRNA in samples collected at different time points were assessed by RNA-seq. Further, a kinetic model with ordinary differential equations was developed that summarizes our understanding of the system. O_2_: oxygen; WC: Wilkins Chalgren medium; BT: *Bacteroides thetaiotaomicron*; RI: *Roseburia intestinalis*.

## Materials and Methods

### Human gut bacterial strains

Bacterial strains of *Bacteroides thetaiotaomicron* VPI-5482 (DSM 2079^T^), DSM 24846, DSM 108160, DSM 108161 and *Roseburia intestinalis* L1-82 (DSM 14610^T^) were obtained from the Deutsche Sammlung von Mikroorganismen und Zellkulturen (DSMZ, Germany). The strains were frozen in Wilkins-Chalgren Anaerobe Broth (WC; Oxoid Ltd., Basingstoke, United Kingdom) plus 20% glycerol and maintained at -80°C until use.

### Mucin bead preparation

Mucin beads were generated based on a protocol previously described [33]. Briefly, a polysaccharide gel was prepared that contained (in g/100 ml) Gelrite gellan gum (2.5, Sigma), xanthan gum (0.25, Sigma) and porcine mucin type II (5, Sigma; pretests of the strains grown in WC medium showed no difference between the beads supplemented with type II and type III mucins). The dried powders were suspended in 100 ml of 90°C pre-heated distilled water containing 0.2 g sodium citrate (Sigma) and 350 μl 10 M NaOH. This polymer solution was mixed until dissolved and then autoclaved for 20 min at 121°C. The bead generation was performed under sterile conditions and was based on a two-phase dispersion process [34]. The autoclaved solution was cooled down to 50°C on a heater and slowly poured with a distance of 40 cm into 45°C sunflower oil while keeping the stirring speed at 300 rpm, to obtain a suspension of aqueous gel droplets in the hydrophobic phase. Gel beads were formed when cooling the suspension down to 15°C on ice. The oil was carefully washed off the beads and were subsequently hardened by soaking in a 0.1 M CaCl_2_ solution (adjusted to pH 5.2) for 20 min at room temperature. Beads with a diameter of 500 μm were selected by wet sieving and stored at 4°C in aseptic conditions until use.

### Cultivation and sample collection

Serum bottles (120 ml) containing 60 ml WC medium and around 8 000 mucin beads were used as a spatially structured system to grow *B. thetaiotaomicron* and *R. intestinalis*. Briefly, each bottle was first filled up with 50 ml WC medium, and 3 g wet mass of 500 μm mucin beads in 10 ml WC were pipetted into each bottle under sterile conditions. The bottles were transferred to the anaerobic workstation (Don Whitley A35 with HEPA filter; 10% H_2_, 10% CO_2_, 80% N_2_, 55% humidity) to be deoxygenated for 120 h. Oxygen-free conditions in liquid and in the beads were confirmed through the color change of redox indicator resazurin. The bottles were closed with rubber stoppers and sealed with aluminum caps that were UV sterilized for 1 h in a laminar flow chamber before use. For bottle experiments without mucin beads, bottles were filled with 60 ml WC medium that was prepared in the same way as described above.

To prepare the inoculum, 100 μl of *B. thetaiotaomicron* or *R. intestinalis* from frozen stocks was first inoculated in a 10-ml serum vial (Wheaton) containing 5 ml WC medium as preculture. These serum vials were prepared by adding autoclaved medium and were deoxygenated in the anaerobic workstation for 24 h. The vials were closed with rubber stoppers and sealed with aluminum caps before use. After 16 h, each species’ preculture was diluted to an OD_600_ of 0.1 determined by a portable spectrometer (DR1900; Hach Lange GmbH, Germany) and used as the inoculum. The serum bottles were inoculated with 1 ml of the diluted preculture and were incubated at a constant stirring rate of 170 rpm at 37°C in a shaker (KS 4000 i; IKA, Staufen, Germany). Cultivations of the monoculture and co-culture were followed for 120 h after inoculation, with samples taken from both the liquid and beads every four hours for the first 48 h and every 12 h afterwards. For further investigation of the *B. thetaiotaomicron* and *R. intestinalis* growth and their interactions in the long term, an experiment was designed for the co-culture to follow for 288 h with samples taken every 24 h since 120 h after inoculation. These batch fermentation experiments were performed in closed bottles, with a controlled environment only at the beginning of an experiment. All experiments were performed at least in three biological replicates and always had a negative control bottle without inoculation but with sampling for each time, to verify its sterility.

For each time point, two milliliters of the fermentation broth containing the suspended mucin beads were sampled to 2-ml tubes (Eppendorf). The serum bottles were opened and closed carefully each time to avoid contamination in the anaerobic workstation. During sampling the beads settled downward to the bottom within seconds. The supernatant was carefully pipetted into new 2-ml tubes in the anaerobic workstation, and subsequently was used to measure pH (PH-922, ISFET, Japan) and OD_600_, and to profile metabolites as well as quantify bacterial cell counts by live/dead staining followed by flow cytometry.

### Spent media and pH preference experiments

Several spent media assays were conducted in serum bottles to clarify how *B. thetaiotaomicron* and *R. intestinalis* interact with each other, and how abiotic factors influenced their growth. First, we grew a culture of *B. thetaiotaomicron* in WC for 48 h without mucin beads and sterilized the spent medium through a filtration with 0.2-μm sterile syringe filters (PES; VWR International). In addition, the filtered medium was autoclaved and deoxygenated for 24 h in the anaerobic workstation. Four conditions were prepared to cultivate *B. thetaiotaomicron* in the spent medium: *i*) the original spent medium without adjusting pH (pH 5) and without adding carbon source, *ii*) spent medium with pH adjusted to 6.5, *iii*) spent medium with glucose and pyruvate added in the same concentrations as in WC at pH 5 and *iv*) spent medium with added glucose and pyruvate plus pH adjusted to 6.5. We grew *B. thetaiotaomicron* in three biological replicates in the serum bottles for a period of 48 h and sampled every 8 h for measuring pH, OD_600_, metabolites and live/dead cell counts. Second, we grew *B. thetaiotaomicron* in WC plus mucin beads in different periods of 24 h, 48 h and 72 h, as well as in WC without mucin beads for 48 h. The cell- and oxygen-free spent media were prepared in the same way as described above, and the pH was adjusted to 6.5 before autoclaving.

We further adjusted the pH of WC medium to the specified values of 4.5, 5.0, 5.5 and 6.0 by using 1 M HCl solution, and also included the original WC of pH 6.7 for testing in serum bottles. The growth of *B. thetaiotaomicron* was followed for 48 h, with samples taken at 0 h, 4 h, 8 h, 12 h, 24 h, 36 h and 48 h and measured as described above.

### Detachment experiment

To estimate the rate at which bacterial cells detach from particles, we followed the population change of planktonic cells in a fresh WC medium, into which mucin beads with attached *B. thetaiotaomicron* cells were added. We first grew *B. thetaiotaomicron* culture in a serum bottle containing 60 ml WC plus mucin beads, with a same inoculum size as described: 1 ml of OD_600_ 0.1. After 24 h cultivation, 2 ml of the fermentation broth containing the suspended mucin beads were sampled to each 2-ml tube. 0.5 ml of phosphate buffer (PBS, pH 7.3; Invitrogen, Thermo Fisher Scientific, USA) was added to each tube after washing with 1 ml PBS. The beads in PBS were transferred to a 12-well plate (Sarstedt, Germany), with 1.5 ml sterile WC medium in each well. For comparison, we included wells of fresh WC medium and WC medium supplemented with sterile mucin beads, both of which were inoculated with *B. thetaiotaomicron* in a volume of 25 µl of OD_600_ 0.1. The experiment was performed in four replicate wells for each group. The plates were incubated at a constant stirring rate of 170 rpm at 37°C, with liquid samples taken at 0 min, 30 min, 60 min, 90 min, 150 min and 240 min for live/dead cell staining followed by flow cytometry.

### Mucin monomeric sugar assays

The degradation products of mucin including *N*-acetylneuraminic acid (Neu5Ac), *N*-acetylgalactosamine, *N*-acetylglucosamine, fucose, mannose and galactose were tested for their effects on the growth of *B. thetaiotaomicron* and *R. intestinalis*. The tests for each species were performed by using 24-well plates (Sarstedt, Germany) filled with two types of media: the standard WC and WC without carbon sources glucose and pyruvate. Each medium was split into eight aliquots, each of which was supplemented with either one of those mucin sugars in 1 mM or a mixture of all six sugars together in 1 mM or without any sugar added as the control. We first inoculated 0.5 ml preculture with OD_600_ 0.1 to 20 ml of each aliquot, and then transferred 1.2 ml to each well in three replicates. The plates of *B. thetaiotaomicron* or *R. intestinalis* were incubated at a constant stirring rate of 120 rpm at 37°C for 24 h or 48 h, respectively, with samples taken every 12 h for pH and live/dead staining followed by flow cytometry measurements.

### Quantification of bacterial cell counts using flow cytometry

DNA-based staining using SYBR Green I (SG; Invitrogen) in combination with propidium iodide (PI; Invitrogen) was applied to quantify and differentiate bacterial cells with intact and damaged cytoplasmic membranes following the protocols described before [35]. Briefly, the stock staining solution contained per ml: 10 μl of SG (10 000 concentrate), 20 μl of 20 mM PI and 970 μl of the filtered dimethylsulfoxide. Cells of the fresh culture in liquid from different time points were diluted in a filter-sterilized PBS buffer for the live/dead cell staining. Dilutions in 1:10 and 1:200 were performed for the first two time points (0 h and 4 h) and the later time points (after 8 h), respectively. Under anoxic conditions, cell samples were diluted in 1-ml 96-well plates (Eppendorf, Germany) and stained with 3 μl of the SG/PI staining solution, and subsequently were incubated for 20 min in the dark at 37°C right before flow cytometric measurements. To quantify the attached cell counts, the mucin beads sampled from different time points were first washed with 1 ml filter-sterilized PBS in 2-ml tubes (Eppendorf). After carefully removing the supernatant, we added another 0.5 ml PBS to the tubes that later were sonicated in the water bath of an ultrasonic cleaner (USC200TH, VWR International) for 5 min at 45 kHz. The fresh detached cells in PBS were diluted 1:10 in 96-well plates for all time points and were used for the live/dead cell staining as described above. To confirm method accuracy, we also used a green fluorescent protein (GFP)-tagged *B. thetaiotaomicron* containing the modified plasmid pWW3452 (CTAATG in the start codon of GFP was replaced with CATATG), which can stably fluoresce when grown under anoxic conditions with subsequent exposure to oxygen for 60 min as described [36].

Flow cytometric counts of the fresh cells were obtained by using a benchtop CytoFLEX S flow cytometer (Beckman Coulter, Brea, USA), equipped with the identical lasers and detectors as described previously [37]. Cell events of each sample were recorded for 1 min at a sample flow rate of 10 μl/min, using the CytExpert software (version 2.4.0.28). Threshold values of 3 000 and 2 000 were applied to the channels of forward scatter and side scatter, respectively. These thresholds were selected based on our tests of *B. thetaiotaomicron* and *R. intestinalis* cells from different growth phases in serum bottles containing WC medium without and with mucin beads to exclude most of the background events and to obtain accurate cell counts. In addition to the instrumental calibration with CytoFLEX Daily QC Fluorospheres, 0.5 μm and 1 μm green fluorescent beads (Thermo Fisher Scientific, USA) were applied to each batch of experiments as internal standards, to monitor the instrument stability and to make cytometric data from different batches more comparable.

### Flow cytometry data processing

We used an in-house pipeline to quantify the absolute abundance of *B. thetaiotaomicron* and *R. intestinalis* cells by clustering populations of flow cytometric events in a UMAP space [38]. The detailed protocol and scripts to reproduce our cell population counts are available at: bit.ly/3WNrslL. Briefly, we normalized and scaled raw flow cytometry data, transforming the values with the arcsin function for uncorrelated unit variances and zero means. We then projected this data into three-dimensional UMAP space using Euclidean distance, with parameters of 25 and 0.1 for the number of neighbors and minimum distance. Flow cytometry runs containing cells and runs containing only the media (blank) allowed classification into four categories: (1) “live” (SG positive, PI negative, and not in the blanks); (2) “inviable” (not in the blanks, but PI positive) and; (3) “debris” (PI and SG negative, but not in the blanks); (4) “blank” (overlap with the blank files). PI and SG positive events were determined based on a threshold signal in their respective channels, which were empirically estimated from the distribution of all the data samples. The overlap with blank events from blank control files was determined by drawing a circle around the points in the blank files, whose radius is proportional to the density of blank points found in its proximity and classifying the area spanned by this circle as blanks in a combined UMAP projection of samples and blank files. This approach, that mimics the process of drawing a circle around the events in the blank file was sufficient because the UMAP projection showed a clear separation between the blank and cell events. Using the density as a basis for the circles allowed us to automatically account for the cases where there is a small carryover from cell events to the blank runs, preventing the automatic detection of cell events as blanks. Only the events classified as “live” were selected for the next steps. We then used the monoculture samples of each respective time point to classify cells in the co-culture samples. For this we first built a training space consisting of a labeled UMAP projection of 5 000 random events selected from each monoculture biological replicate. Next, we used supervised UMAP to reproject the monoculture and co-culture live-cell events into the training space. We then used the *K*-nearest neighbors vote classifier (with parameters: n_neighbors = 50, weights = distance, and metric = Mahalanobis) to identify the species labels of the co-culture events based on their proximity to the monoculture events in the supervised UMAP space. Before filtering out the blanks, each sample was overlayed with their corresponding blank runs and manually inspected using clickable three-dimensional scatter plots, where we could visually assure that the populations classified as blank in our samples overlap with the populations in the blank files and not with the cell populations. Examples of these scatter plots are available at: bit.ly/3WNrslL. Similarly, we manually verified that each classified species population overlapped with their monoculture runs for similar time points. The manual inspection also showed that our parameter choices for UMAP and the *K*-nearest vote classifier were robust.

### Contamination check and confirmation of abundance profiles with 16S rRNA genes

Sanger sequencing of 16S rRNA genes was performed regularly to check each strain identity. In addition, we selected four biological replicates for each time point from the co-cultivation experiments of *B. thetaiotaomicron* and *R. intestinalis* in WC with and without mucin beads, to confirm the abundance profile of *B. thetaiotaomicron* and *R. intestinalis* in co-culture with 16S rRNA gene amplicon sequencing (Figure S1). Sequencing was performed using the MiSeq platform (Illumina) to generate paired-end reads of 250 base pairs. The taxonomy was assigned by using the RDP classifier v2.13 with a confidence threshold of 80% [39].

### Metabolite profiling

Liquid samples in 1 ml were collected and centrifuged for 20 min at 21 130 × *g* at 4°C (Centrifuge 5424 R; Eppendorf, Hamburg, Germany). The supernatants were used for chemical analyses by applying external standards for calibration and quality control. For the calibration curves, all compounds (Merck) were in high purity and acidic form (except sugars), and were dissolved or diluted in ultrapure water for calibration. The concentrations of trehalose, glucose, pyruvate, succinate, formate, acetate, propionate, lactate, *n*-butyrate and Neu5Ac were determined by high-performance liquid chromatography (HPLC). HPLC analysis was carried out on an Agilent 1200 series HPLC system (Agilent Technologies, USA) equipped with a Bio-Rad Aminex HPX-87H column (Bio-Rad, USA) and a refractive index detector at 40°C. Samples in 20 μl were injected and eluted for 60 min using 5 mM H_2_SO_4_ as the mobile phase at a flow rate of 0.6 ml/min.

Concentrations of mucin monomeric sugars galactose, fucose and mannose were relatively quantified by using gas chromatography, carried out by the Metabolomics Core of KU Leuven. The supernatant in 100 μl of each sample was diluted to 1 ml in 80% methanol, and left overnight to allow the precipitation of lipids and proteins. After centrifugation, 100 μl of the supernatant was transferred and evaporated to dryness on a speedvac. A 2% methoxyamine solution was prepared by dissolving methoxyamine hydrochloride (Supelco, ref. 226904-5g) in pyridine (Sigma-Aldrich, ref. 270970-1L). 25 μl of the methoxyamine solution was added to each sample and left to react for 1.5 h at 37°C for methoximation of the hexoses. Afterwards, 80 μl N,O-bis(trimethylsilyl)trifluoroacetamide with 1% trimethylchlorosilane (Supelco, ref. 15238) was added and left to react for 30 min at 60°C for trimethylsilylation of the methoximated hexoses. One microliter of each methoximated and trimethylsilylated sample was injected by 2:1 split injection by an Agilent Technologies 7693 autosampler (Agilent Technologies, Santa Clara; California) into an Agilent 7890A gas chromatograph equipped with a fused silica capillary column (0.25 mm × 30 m) with a chemically bonded 0.25 μm HP-5MS stationary phase (Agilent J&W GC columns, Folsom; California, ref 19091S-433). The injector temperature was 250°C, and the septum purge flowrate was 3 ml/min. The helium flowrate through the column was 0.9 ml/min. The column temperature was held at 160°C for 23.5 min and then increased by 100°C/min to 300°C, with 1 min holding there. The column effluent then entered an Agilent 7000 triple quadrupole mass spectrometer. Temperatures of the transfer line and electron impact ion source were 250°C and 230°C, respectively. Ions were generated by a 70 eV electron beam. Spectra were acquired in a selected ion monitoring mode, with fragment masses based on the analyses of standards.

### RNA extraction and sequencing

We included a total of 69 samples representing the different growth phases of *B. thetaiotaomicron* and *R. intestinalis* in three biological replicates for RNA sequencing, which were selected from the following six independent experiments: monoculture *B. thetaiotaomicron* in WC (4 h, 12 h and 36 h), monoculture *R. intestinalis* in WC (4 h, 12 h and 48 h), coculture *B. thetaiotaomicron* and *R. intestinalis* in WC (12 h, 24 h and 36 h), monoculture *B. thetaiotaomicron* in WC plus mucin beads (4 h, 12 h, 24 h, 48 h and 120 h), monoculture *R. intestinalis* in WC plus mucin beads (16 h, 28 h, 40 h and 48 h), and coculture *B. thetaiotaomicron* and *R. intestinalis* in WC plus mucin beads (6 h, 12 h, 24 h, 48 h and 120 h).

Total RNA was extracted and purified from RNAprotect-treated (Qiagen, Hilden, Germany) frozen liquid samples using the phenol-free RNeasy Plus Mini Kit (Qiagen) according to the instructions of the manufacturer. The cell pellets from -80°C were first lysed by adding the lysis buffer RLT from the kit with 10 μl/ml β-mercaptoethanol. The content of PowerBead tube (glass, 0.1 mm; Qiagen) was further added to the lysate prior to using a bead beater (TissueLyser II, Qiagen) for homogenizing the mixture for 5 min at a speed of 30 beats/sec. Other steps were performed by following the instructions. The eluted RNA samples in nuclease free water were stored at -80°C until use. RNA concentrations were determined with a Nanodrop ND 1000 spectrophotometer (Thermo Fisher Scientific, USA) and a Qubit 2.0 fluorometer using the Qubit RNA High Sensitivity Assay Kit (Invitrogen). Samples with a 260/230 absorbance ratio below 0.5 were purified using the MinElute PCR Purification kit (Qiagen) and re-measured afterwards, with all samples above 0.5 after purification. RNA integrity and yield were evaluated using RNA Nano and Pico 6000 LapChips (Agilent Technologies, Santa Clara, CA, USA) run on an Agilent 2100 Bioanalyzer (Agilent Technologies).

Library preparation and sequencing of RNA-seq libraries were carried out by the Genomics Core of KU Leuven. All samples were prepared using the NEBNext Ultra II Directional RNA Library Prep Kit for Illumina (New England BioLabs, #E7760L), with an initial amount of 100 ng total RNA in 12 μl. Bacterial rRNA depletion was proceeded using the rRNA Depletion Kit (Bacteria) (New England BioLabs, #E7850L), and the residual mRNA was quality controlled and converted to the libraries that were sequenced on a NovaSeq 6000 (Illumina, San Diego, CA, USA) on a v1.5 S4_300 kit in XP mode. The resulting data of paired-end reads in fastq file format were obtained and demultiplexed before further analysis.

### RNA-seq data processing

The analysis of the raw sequencing reads was performed as follows: low quality reads and adapters were trimmed with fastp [40]. High quality RNA reads were mapped to their respective reference transcripts with Salmon run in a selective alignment mode and with a decoy-aware index constructed from each organism’s genome [41]. The latest reference genomes and transcripts for *B. thetaiotaomicron* and *R. intestinalis* were downloaded from the ensemble bacteria [42] and BV-BRC databases [43], respectively: GCA_000011065.ASM1106v1.dna.toplevel.fa (*B. thetaiotaomicron* VPI-5482), GCA_000011065.ASM1106v1.cds.all.fa (*B. thetaiotaomicron* VPI-5482), GCA_900537995.1_Roseburia_intestinalis_strain_L1-82_genomic.fna (*R. intestinalis* L1-82), GCA_900537995.1_Roseburia_intestinalis_strain_L1-82_cds_from_genomic.fna (*R. intestinalis* L1-82). Raw transcript counts were extracted from salmon’s output and used for differential gene expression analysis. COG categories, COG numbers and KO numbers were extracted from NCBI COG and KEGG database [44, 45], for *B. thetaiotaomicron*. To obtain annotations for *R. intestinalis*, we mapped the respective transcripts to the eggNOG database v5.0 with the eggNOG-mapper 2.1.9 [46, 47].

### Statistics

To measure gene expression levels, we used methods of transcripts per million and DESeq2’s median of ratios to normalize RNA read counts [48]. A strict cutoff of at least 1.5-fold change and a Benjamini-Hochberg adjusted *p* value less than or equal to 0.01 were applied for differential gene expression calculated by DESeq2. Other statistical analyses were performed using R (version 4.2.0).

### Modelling

Details of model definition and parameterization can be found in the Supplementary file 2.

## Results

### Different responses to sugar depletion

Glucose and pyruvate were provided as the main carbon sources in WC medium (Figure 2a). When grown in monoculture, *B. thetaiotaomicron* produced succinate, lactate, formate, and acetate during fermentation, which decreased the pH from 6.7 to 5.0. At 20 h, *B. thetaiotaomicron* reached its maximal density of 1.2 × 10^6^ viable cells/µl and hereafter its density decreased. After the depletion of glucose and pyruvate, the acidic conditions led to a loss of viable *B. thetaiotaomicron* cells but did not affect the total cell count. Loss of viability at low pH (5.0) was further confirmed by spent media and pH experiments. We re-grew *B. thetaiotaomicron* in its spent medium (collected at 48 h cultivation when glucose is depleted), and found that an initial pH of 6.5 is required for growth (on trehalose; Figure S2). Supplementing the spent medium with glucose and pyruvate only enhanced growth when the pH was also increased (Figure S2a). In addition, *B. thetaiotaomicron* did not grow on glucose and pyruvate when the pH of WC medium was lowered to 4.5 and 5.0, respectively, and its growth rate was decreased at pH 5.5 and 6.0 compared to pH 6.7 (Figure S2b). *R. intestinalis* in monoculture produced lactate, formate, acetate and butyrate from glucose and pyruvate, corresponding to a pH decrease from 6.7 to 5.8 at 24 h and reached a maximal density of 7.8 × 10^5^ viable cells/µl (Figure 2b). After the depletion of glucose and pyruvate in 24 h, *R. intestinalis* gradually entered a slow growth mode, seen as a drop in viable cells to 1.8-3.8 × 10^5^ cells/µl from 40 h to 120 h in two independent experiments. In addition, we observed a further significant (*p* < 0.001 by Student’s t test) increase of butyrate concentration concurrent with decreasing acetate and, in some cases, lactate (Figure 2b and Figure S3 for lactate consumption in *R. intestinalis* monoculture), resulting in a pH increase from 5.7 to 6.0. In brief, viable *B. thetaiotaomicron* cells decline primarily due to the lowered pH, whereas *R. intestinalis* maintains higher cell densities by entering a slow growth mode.

**Figure 2.**
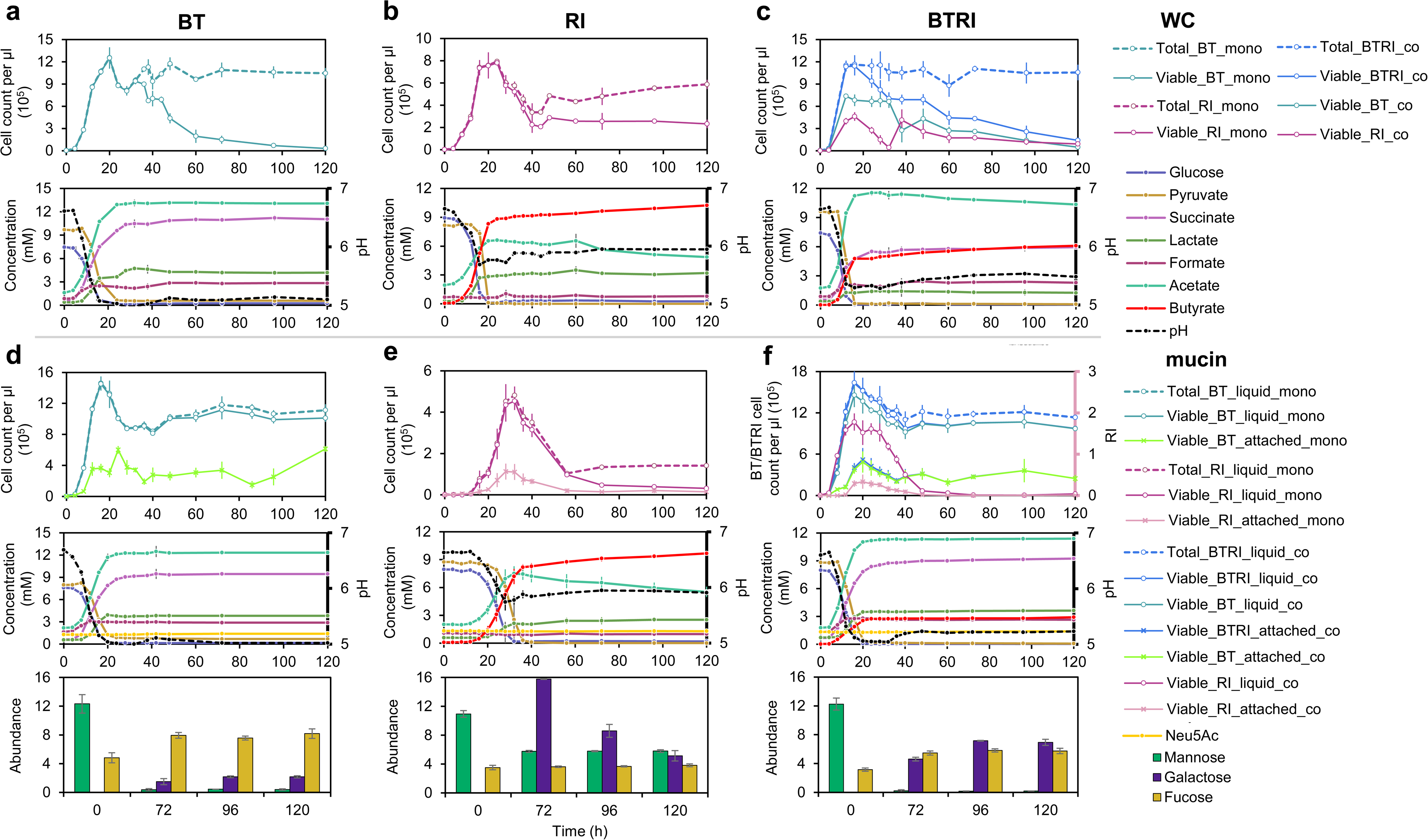
Performance of *B. thetaiotaomicron* and *R. intestinalis* in monoculture and co-culture without and with mucin beads during batch fermentation. Cell density, concentrations of glucose, pyruvate and fermentation products, and pH values for *B. thetaiotaomicron* monoculture **(a)**, *R. intestinalis* monoculture **(b)** and their co-culture **(c)** in WC medium during 120 h cultivation; Cell density, concentrations of glucose, pyruvate and fermentation products, pH values as well as concentrations of mucin monomeric sugars for *B. thetaiotaomicron* monoculture **(d)**, *R. intestinalis* monoculture **(e)** and their co-culture **(f)** in WC with mucin beads during 120 h cultivation. Circles represent the mean values of biological replicates. Error bars indicate the standard deviation of biological replicates (n=5 for *B. thetaiotaomicron* and *R. intestinalis* monocultures in WC experiment; n=6 for co-culture in WC; n=3 and n=5 for *B. thetaiotaomicron* and *R. intestinalis* monocultures in WC with mucin beads, respectively; n=5 for co-culture in WC with mucin beads). Concentrations of mucin sugars are represented by the integrated area under the peak corresponding to the metabolite. To show the same range on the y axis of barplots, we divided these values by factors of 1 000 000, 5 000 000 and 10 000 for galactose, mannose and fucose, respectively. WC: Wilkins Chalgren medium; BT: *Bacteroides thetaiotaomicron;* RI: *Roseburia intestinalis*; mono: monoculture; co: co-culture; Neu5Ac: *N*-acetylneuraminic acid.

Given the observed diauxic shift from glucose/pyruvate to lactate/acetate, we performed RNA-seq of *R. intestinalis* grown in monoculture for several time points and replicates to investigate its metabolic mechanism. During the exponential growth period, genes associated with glycolytic and pyruvate-utilizing pathways were more highly expressed at mid-log phase of 12 h compared to at 4 h (Figure S4b). Although glucose ran out at 48 h, we found that *R. intestinalis* kept expressing glycolysis genes and also non-essential genes such as those involved in cell motility despite growing in a well-mixed environment (Figure S5). We further compared the effect of different normalization methods (transcripts per million and DESeq2’s median of ratios [48]), which showed consistent results except for some glycolysis genes (Figure S6). When comparing the transcript levels of *R. intestinalis* in monoculture between 12 h and 48 h, the significant up-regulation of genes encoding lactate dehydrogenase (RIL182_00335 and 01719) and butyryl-CoA:acetate CoA-transferase (RIL182_00413) corroborates the finding that *R. intestinalis* converts lactate/acetate to butyrate after glucose depletion. The energy yielded by butyrate production from lactate/acetate (Δ_f_*G*^0^ = -46 KJ/mol) is much lower than that from glucose (Δ_f_*G*^0^ = -250 KJ/mol) [49]. Therefore, it is not surprising that *R. intestinalis* grows more slowly on lactate/acetate. In summary, *R. intestinalis* maintains its slow growth in the absence of glucose by converting acetate and lactate to butyrate.

### Effects of mucin on monoculture

*B. thetaiotaomicron* survives longer in the presence of mucin beads than without them. After depleting glucose and pyruvate at 24 h, planktonic *B. thetaiotaomicron* retained a viable cell count higher than 1.0 × 10^6^ cells/µl (Figure 2d), which was also maintained when further extending the cultivation period from 120 h to 288 h (Figure S7a; Figure S7b for three non-type strains). In contrast, the growth of *R. intestinalis* on glucose/pyruvate seems to be impeded by mucin, demonstrated by significantly lower cell densities at the same time points and lower maximum of 4.5 × 10^5^ viable cells/µl at 36 h when compared with *R. intestinalis* grown without mucin beads (Figure 2e; Figure S8). After nutrient depletion, *R. intestinalis* still exhibited the slow growth mode characterized by a further increase of butyrate and decrease of acetate, but with a much lower density of 0.3-0.5 × 10^5^ viable cells/µl than in the absence of mucin. In addition to the planktonic cells in liquid, we also quantified the cells of *B. thetaiotaomicron* and *R. intestinalis* attached to the mucin beads at different time points. Attached *B. thetaiotaomicron* and *R. intestinalis* in monoculture reached maximal densities of 6.0 × 10^5^ viable cells/µl at 24 h and 1.1 × 10^5^ viable cells/µl at 32 h, respectively (Figure 2d-e). Thus, in the presence of mucin, *B. thetaiotaomicron* can maintain high densities in the long term, whereas the density of *R. intestinalis* declines.

Regarding the mucin degradation products, the concentration of Neu5Ac showed a significant (*p* < 0.001 by Student’s t test) increase of 0.12 mM in *B. thetaiotaomicron* monoculture after 120 h and no significant change in *R. intestinalis* monoculture (Figure 2d-e). We also observed higher levels of fucose and galactose for *B. thetaiotaomicron* grown with mucin beads than without it. The dynamics of galactose in *R. intestinalis* monoculture indicates that *R. intestinalis* also hydrolyzed mucin and used resulting degradation products such as galactose. To investigate the effects of mucin degradation products on the growth of *B. thetaiotaomicron* and *R. intestinalis* at different glucose levels, monosaccharides including Neu5Ac, *N*-acetylgalactosamine, *N*-acetylglucosamine, fucose, galactose, mannose, and their mixture were added to the standard WC medium and WC medium without glucose and pyruvate in 24-well plates. In the presence of glucose/pyruvate, none of the supplemented sugars influenced *B. thetaiotaomicron* growth significantly (*p* > 0.05 by Student’s t test) compared to the WC control (Figure S9a). In contrast, Neu5Ac increased *R. intestinalis* growth by 43% while other sugars had negative effects (14%-46%; Figure S9c). In the absence of glucose/pyruvate, all sugars except Neu5Ac had positive effects on *B. thetaiotaomicron* growth (2%-35%; Figure S9b), and all sugars, in particular galactose, promoted *R. intestinalis* growth (13%-197%; Figure S9d). In conclusion, mucin-derived monosaccharides have species-specific effects that in addition depend on the level of glucose.

The RNA-seq data further elucidated the different responses of *B. thetaiotaomicron* and *R. intestinalis* to mucin. For *B. thetaiotaomicron*, we first collected a list of genes/operons associated with the polysaccharide utilization loci of mucin *O*-glycans, which were possibly activated by two types of transcriptional regulators, namely extracytoplasmic function sigma/anti-sigma pair (ECF-σ) and hybrid two-component system (HTCS) [50]. The genes linked to HTCS and ECF-σ showed significantly high expression in the first 4 h (Figure 3a; *p* < 0.001 for both by Student’s t test), and genes of ECF-σ-linked loci were also expressed after the glucose/pyruvate depletion (48 h and 120 h), indicating BT utilization of mucin *O*-glycan components. Specifically, after glucose/pyruvate ran out at 24 h, we observed a significantly (*p* < 0.001 in all cases; Student’s t test) increased expression of genes encoding the extracellular galactosidase (BT_3340, BT_4684) and fucosidase (BT_1842), as well as genes involved in the utilization of galactose (BT_0370-71, BT_0623) and fucose (BT_3606, BT_1273-75). These results are consistent with *B. thetaiotaomicron* metabolizing mucin *O*-glycans including galactose and fucose (Figures 2d and S9b). In addition, *R. intestinalis* consistently increased the expression of genes involved in glycolysis, pyruvate utilization, mucin hydrolysis, galactose and mannose utilization, and the butyrate-producing process after depletion of glucose/pyruvate, with or without mucin beads present in the medium (Figures 3b and S4b). At a high glucose/pyruvate level (Figure S10a), the presence of mucin significantly reduced the expression of genes associated with the transport of glucose (RIL182_03793, RIL182_03794, RIL182_03796), pyruvate (RIL182_01689), galactose (RIL182_01067-69) and *N*-acetylglucosamine (RIL182_02688-89) across the outer membrane. At a low glucose/pyruvate level, genes of lactate dehydrogenase (RIL182_00335) and butyryl-CoA:acetate CoA-transferase (RIL182_00413) were significantly down-regulated (Figure S10b). Thus, mucin supports the long-term survival of *B. thetaiotaomicron*, while it undermines *R. intestinalis*’ growth on glucose/pyruvate and possibly also on lactate/acetate.

**Figure 3.**
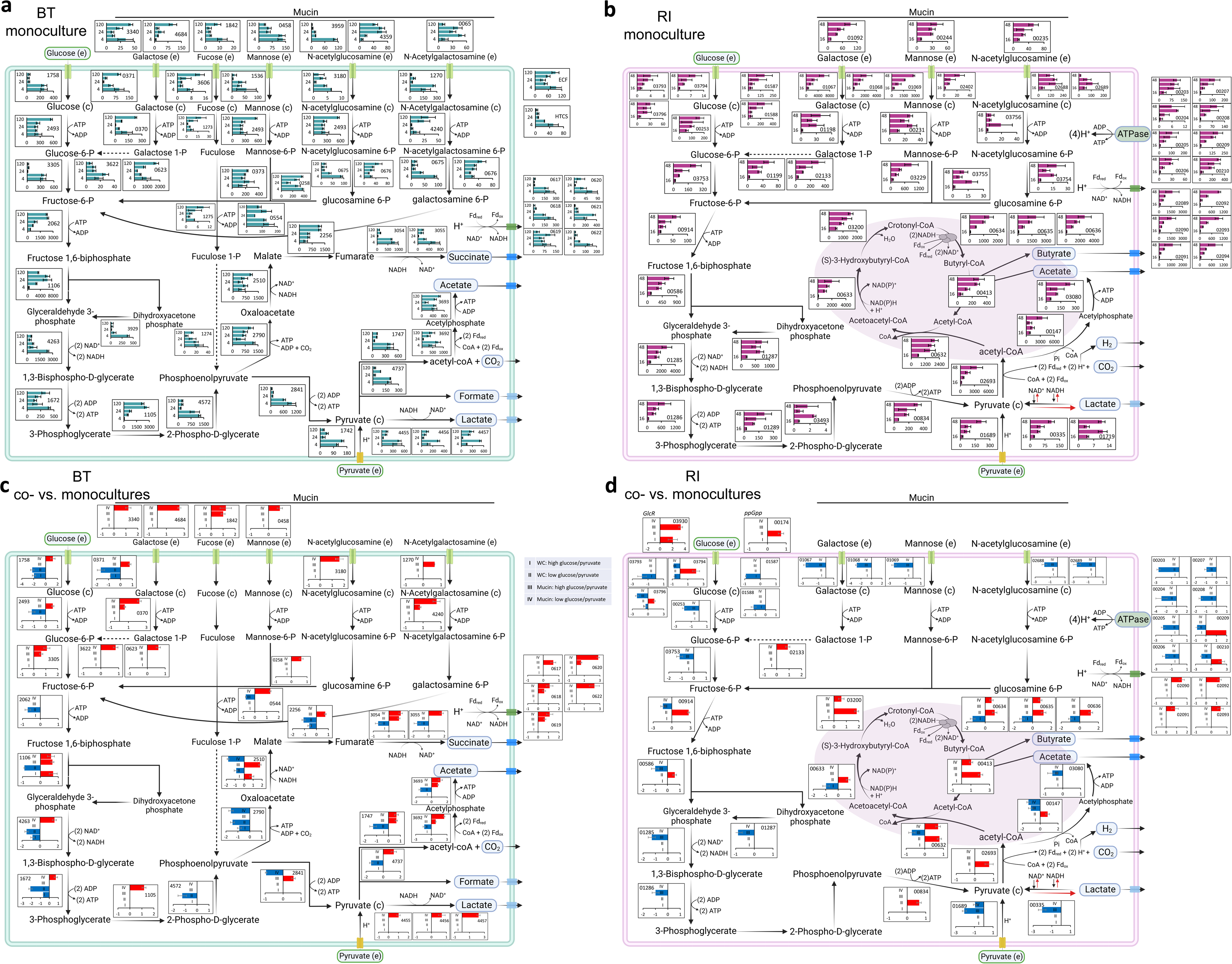
Metabolic maps summarizing gene expression in monocultures and significantly differential expression in co-versus monocultures of *B. thetaiotaomicron* and *R. intestinalis*. Gene expression of the key metabolic routes of glucose and pyruvate fermentation, as well as mucin hydrolysis and mucin sugars utilization in *B. thetaiotaomicron* (a) and *R. intestinalis* (b) monocultures in WC medium with mucin beads; differential gene expression between co-culture and monocultures of *B. thetaiotaomicron* (c) and *R. intestinalis* (d). In each barplot of **a** and **b**, gene expression levels (x axis) were measured in transcripts per million of RNA samples at time (y axis) 4 h, 12 h, 24 h, 48 h and 120 h for *B. thetaiotaomicron*, and at time 16 h, 28 h, 40 h and 48 h for *R. intestinalis*. In each barplot of **c** and **d**, differential gene expression was calculated with DESeq2 for four categories: high levels of glucose/pyruvate in WC (I), low levels of glucose/pyruvate in WC (II), high levels of glucose/pyruvate in WC with mucin beads (III) and low levels of glucose/pyruvate in WC with mucin beads (IV); positive and negative values of log_2_fold-change are shown in red and blue on the x axis, respectively. Numbers in each barplot stand for the Gene ID in the genomes of *B. thetaiotaomicron* and *R. intestinalis*. The solid and dashed lines represent single- and multi-enzyme reactions between the two indicated molecules, respectively. Error bars indicate the standard deviation of three biological replicates. Metabolic maps with gene expression in *B. thetaiotaomicron* and *R. intestinalis* monocultures in WC without mucin beads are shown in Figure S4 and the Supplementary files 3 and 4. ECF: extracytoplasmic function sigma/anti-sigma pair; HTCS: hybrid two-component system; GlcR: HTH-type transcriptional repressor of glucose; ppGpp: alarmone guanosine-5’-3’-bispyrophosphate.

### Impact of mucin on interaction dynamics

For the co-culture grown in WC, the co-utilization of glucose and pyruvate resulted in two separate density peaks of 7.4 and 4.6 × 10^5^ viable cells/µl for *B. thetaiotaomicron* and *R. intestinalis*, respectively (Figure 2c). The pH decreased from 6.7 to 5.3, which was not as acidic as in the *B. thetaiotaomicron* monoculture (5.0) but lower than in *R. intestinalis* monoculture (5.7). After depletion of glucose and pyruvate, *B. thetaiotaomicron*’s viable cell numbers gradually declined, as in monoculture. *R. intestinalis* presented a second peak of 4.1 × 10^5^ viable cells/µl at 38 h. In addition, the net concentration of lactate in co-culture was significantly lower than in each monoculture (*p* < 0.001 in both cases; Student’s t test). For the co-culture grown in the presence of mucin beads, *B. thetaiotaomicron* reached a two-fold higher maximal density than without beads (1.5 × 10^6^ viable cells/µl at 16 h). In contrast, *R. intestinalis* in the mucin co-culture exhibited significantly lower cell densities than in WC co-culture (*p* < 0.01 by Student’s t test), accompanied by a lower maximal density of 1.8 × 10^5^ viable cells/µl. The pH of 5.1 was nearly as acidic as in the *B. thetaiotaomicron* monoculture with or without mucin. A second peak of *R. intestinalis* density in mucin co-culture was observed at 120 h, corresponding to a roughly ten-fold increase in density from 96 h to 120 h (Figure S7a). The significant (*p* < 0.001 by Student’s t test) increase of galactose in the co-culture compared to the *B. thetaiotaomicron* monoculture suggests that both species hydrolyzed mucin (Figure 2f).

To investigate the interaction dynamics of *B. thetaiotaomicron* and *R. intestinalis* and mucin’s impact on it, we compared viable cell counts in co-culture and monoculture over time. We computed the interaction strength, here defined as the impact of one organism on the other’s abundance, as the log ratio of viable cell counts in liquid in co- versus monocultures. The interaction strength and in some cases its sign (i.e., positive or negative impact) changed over time for both organisms with and without mucin beads (Figure S11). In the presence of glucose/pyruvate, the competition for these nutrients resulted in negative interaction strengths for both organisms. After depletion of glucose/pyruvate and in the absence of mucin, *R. intestinalis* impacts *B. thetaiotaomicron* positively, likely by increasing the pH in the co-culture compared to the *B. thetaiotaomicron* monoculture. In the presence of mucin, *B. thetaiotaomicron*’s impact on *R. intestinalis* during the initial growth on glucose/pyruvate (20-48 h) changed from negative to positive, likely by reducing the concentration of mucin degradation products that affect *R. intestinalis*’ growth negatively in the presence of glucose. In summary, changing conditions alter interaction mechanisms between the two species, which lead to varying interaction signs and strengths.

To further elucidate their changing interaction mechanisms across growth phases, we compared the differential gene expression between co-culture and monoculture (Figure 3c-d). In the presence of *R. intestinalis*, *B. thetaiotaomicron* genes associated with the transport of glucose (BT_0371, BT_1758) were significantly down-regulated in WC without mucin (Figure 3c; I, II). In addition, the expression of six out of ten glycolysis genes as well as genes involved in succinate, acetate and formate production was significantly lowered in WC at low glucose/pyruvate levels (Figure 3c; II). With mucin, *B. thetaiotaomicron* up-regulated glycolysis genes and genes involved in producing formate, acetate, succinate and lactate (Figure 3c; III, IV). At low nutrient levels, it also up-regulated genes encoding the exocellular galactosidase (BT_3340, BT_4684), fucosidase (BT_1842), mannosidase (BT_0458) and sialidase (BT_0455) in response to mucin hydrolysis, and genes associated with the utilization of galactose and *N*-acetylgalactosamine (Figure 3c; IV). *B. thetaiotaomicron* genes (BT_0617-22) belonging to the membrane-associated, proton-translocating ferredoxin:NAD^+^ oxidoreductase (*Rnf* complex) were also more expressed with mucin (Figure 3c; III, IV). In conclusion, *B. thetaiotaomicron* down-regulates the genes of glycolysis and pyruvate oxidation when *R. intestinalis* is present in WC but does the opposite when mucin is also present.

In the presence of *B. thetaiotaomicron*, *R. intestinalis* genes associated with glucose transport (RIL182_03793-94, RIL182_03796, RIL182_01587-88) and the ATP generation via proton pump (RIL182_00203-10) were down-regulated at high and low glucose/pyruvate levels, respectively (Figure 3d; I and II, respectively). A candidate gene of the alarmone ppGpp biosynthesis was significantly up-regulated (RIL182_00174). It is part of *R. intestinalis*’ response to stress such as carbon starvation and may be involved in triggering its diauxic shift. *R. intestinalis* genes associated with the import of glucose (RIL182_03794, RIL182_03796), pyruvate (RIL182_01689), galactose (RIL182_01067-69) and *N*-acetylglucosamine (RIL182_02688-89), as well as seven out of ten glycolysis genes were systematically down-regulated at high glucose/pyruvate levels with mucin (Figure 3d; III). In addition, *R. intestinalis* genes associated with the butyrate-producing process via butyryl-CoA:acetate CoA-transferase pathway were more expressed when glucose/pyruvate was depleted, with or without mucin (Figure 3d; II, IV). Thus, *B. thetaiotaomicron* reduces *R. intestinalis*’ import of glucose and may assist its production of butyrate from lactate/acetate, regardless of mucin supplementation.

### Kinetic model reproduces the observed dynamics

To reproduce physiological observables such as total and viable cell counts as well as concentrations of key nutrients and metabolites, we developed a model with ordinary differential equations that summarizes our understanding of the interaction mechanisms (Figure 4a). Since we see a roughly linear relationship between counts of viable attached and free-swimming cells (Figure S12), the model assumes that a fixed fraction of the cells attaches to mucin beads and that only these cells can degrade mucin. Mucin is treated as an inexhaustible resource. We further assume that changes in SCFAs fully determine changes in the pH, since we observed a strong linear relationship between pH and the sum of the concentrations of SCFAs (Figure S13). Based on the literature, we also assume that a low pH affects the growth of *B. thetaiotaomicron* more strongly than that of *R. intestinalis* (i.e., *B. thetaiotaomicron* is more sensitive to acidity [51]). In addition, *B. thetaiotaomicron* and *R. intestinalis* can both grow on glucose and mucin sugars, which was well-supported by our fermentation data and also confirmed at the transcriptional level. Given our microtiter plate data, we further assume that mucin degradation products inhibit *R. intestinalis*’ growth on glucose (Figure S9c). *R. intestinalis* only switches to its lactate/acetate-consuming slow growth mode when the concentrations of glucose and mucin degradation products are low (Figure S10b). Finally, we hypothesize that glucose inhibits *B. thetaiotaomicron*’s growth on mucin sugars, which is supported by the significant overexpression of the master regulator of carbohydrate utilization in the presence of glucose (log_2_fold-change of BT_4338 in mucin monoculture: 1.8 for 4 h vs 12 h, 3.5 for 4 h vs 24 h and 1.6 for 12 h vs 24 h). We simplified the model by treating glucose and pyruvate, acetate and lactate as well as mucin sugars as single entities (see model scheme in Figure 4a). The slow growth mode of *R. intestinalis* and the behavior of *B. thetaiotaomicron* on mucin after depletion of sugar are represented by separate states (in both cases referred to as “slow growth mode”). Parameter values including metabolite production and consumption as well as state switching rates were determined manually. The equations, parameter values and initial conditions are given in the Supplementary file 2.

**Figure 4.**
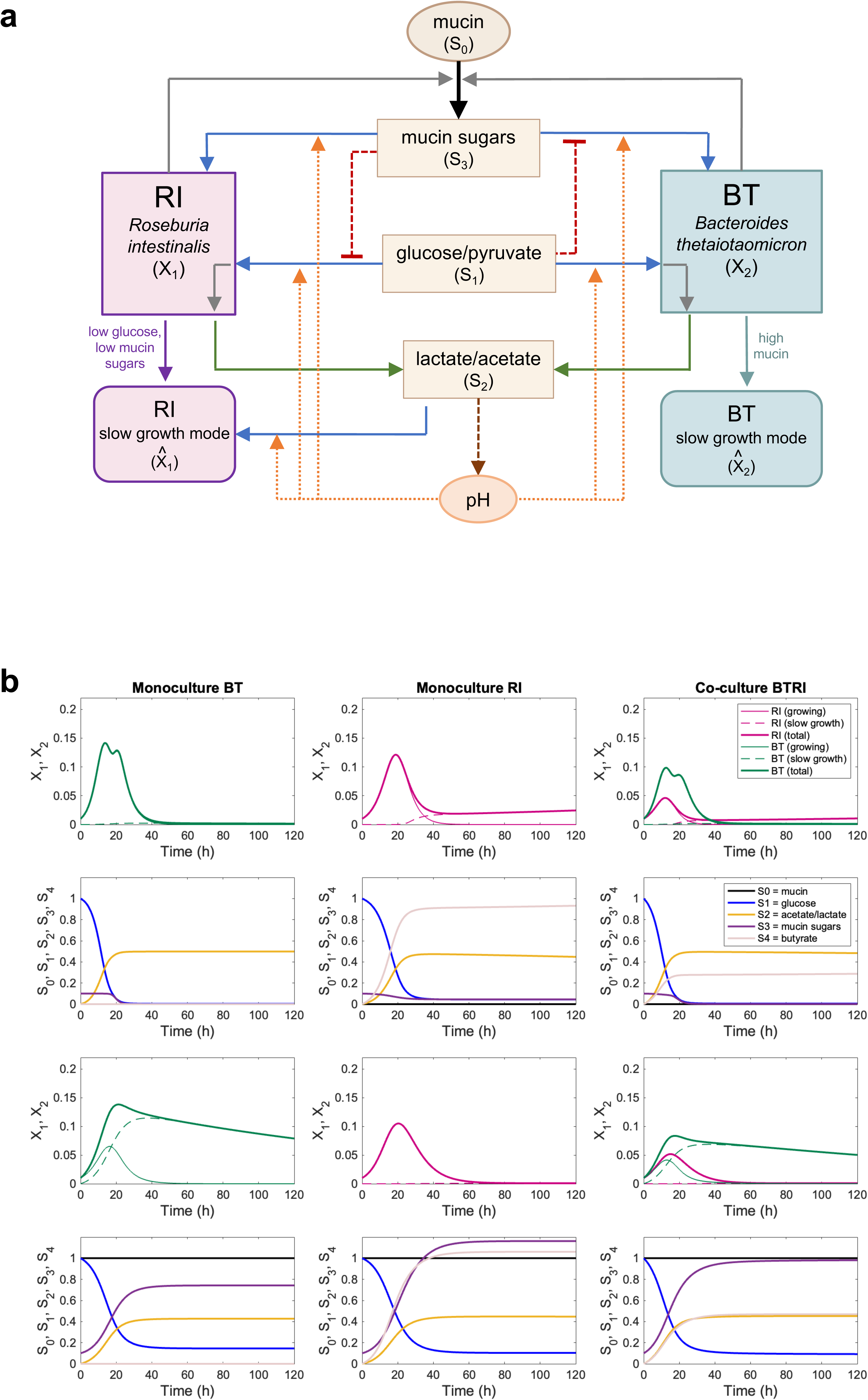
Model scheme and simulations reproducing observations qualitatively. **a**, Scheme of the kinetic model. **b**, Dynamics generated by the model without and with mucin. The equations, parameter values and initial conditions are given in Supplementary file 2.

The model qualitatively reproduces our observations, including the second peak observed for *B. thetaiotaomicron* at 32 h in WC monoculture, the long-term survival of *R. intestinalis* in WC and of *B. thetaiotaomicron* in the presence of mucin beads (Figure 4b). However, the model fails to reproduce the second peak for *R. intestinalis* in WC in co-culture, implying that additional mechanisms are at play that are not yet fully understood.

## Discussion

Here, we investigated the molecular mechanisms underlying the ecological interactions between two representative gut bacteria in the presence of simple and complex carbon sources within a spatially structured environment (Figure 5). The greater affinity of *B. thetaiotaomicron* for the mucin beads (with a mean percentage of 33% attached cells out of total cells, compared to 25% for *R. intestinalis*), coupled with the more beneficial effect of mucin on the growth of *B. thetaiotaomicron* than *R. intestinalis*, agrees with their known niches in the human gut: *B. thetaiotaomicron* thrives as a mucus resident, while *R. intestinalis* tends to inhabit the lumen [17, 32].

**Figure 5.**
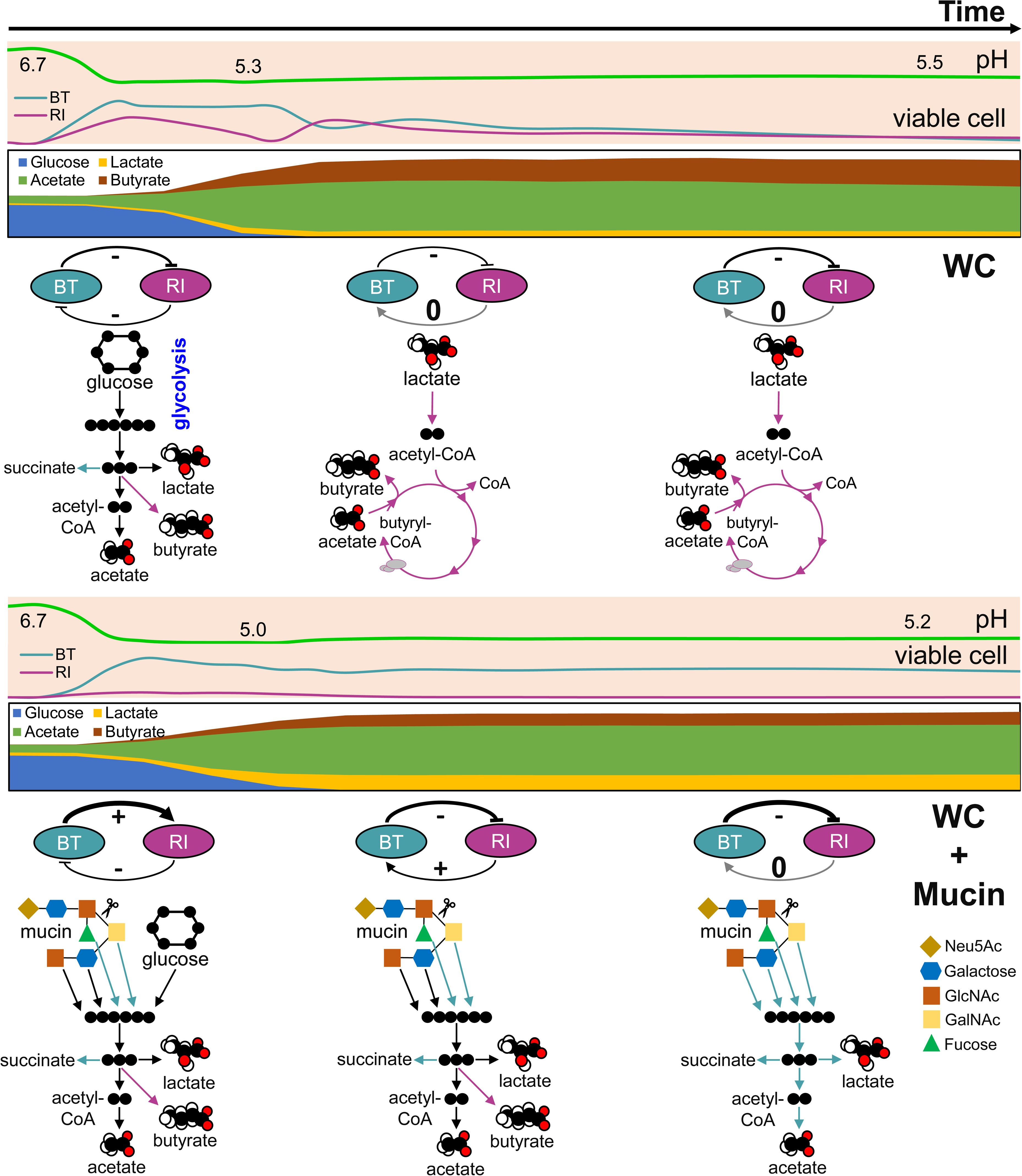
Summary of the interaction mechanisms driving abundances and metabolite concentrations in the *B. thetaiotaomicron* and *R. intestinalis* co-cultures without and with mucin beads. Signs of positive, negative and absence of an interaction are shown as +, - and 0, respectively. The thickness of the arrow represents the interaction strength quantified as the logarithm of viable cell density ratios in co- versus monocultures (detailed in Figures S8 and S11). Main metabolic pathways including glycolysis, converting lactate/acetate to butyrate, mucin hydrolysis and mucin sugars utilization are colored in teal, pink and black for *B. thetaiotaomicron*, *R. intestinalis* and both, respectively. Except for the metabolic pathways inferred from RNA-seq data, others shown in the figure are representations of real data. Neu5Ac: *N*-acetylneuraminic acid; GlcNAc: *N*-acetylglucosamine; GalNAc: *N*-acetylgalactosamine.

We found that even when feeding on similar resources, *B. thetaiotaomicron* and *R. intestinalis* used alternative strategies to cope with and anticipate nutrient scarcity. Growth data combined with temporal gene expression patterns suggest that *B. thetaiotaomicron* acts as a ”funnel” with its central carbon/energy metabolism capable of tapping several monosaccharides. Independent entry points are funneled into glycolysis until phosphoenolpyruvate is produced, which is converted into oxaloacetate by fixing a CO_2_ molecule with the generation of an extra ATP [52]. CO_2_ could be supplied from either the external environment or the conversion of pyruvate to acetyl-CoA. Although this strategy results in accelerated growth, it comes at the cost of significantly inferior performance when actively consumed nutrients are depleted. In contrast, *R. intestinalis* uses carbon chain elongation to generate additional energy and maintain a redox balance (Figure 3b), resulting in butyrate production. Although its growth is slower than that of *B. thetaiotaomicron*, it can switch its energy source to acetate and lactate, which are also produced by *B. thetaiotaomicron*. This diauxic shift resembles the nutrient utilization strategies adopted by other bacterial species [53], mostly well characterized in *E. coli* [54–57]. Some butyrate producers such as *R. intestinalis* can overcome the energetic barrier to convert lactate to pyruvate by employing the flavin-based electron confurcation: driving electron flow from lactate to NAD^+^ at the cost of exergonic electron flow from reduced ferredoxin to NAD^+^. However, producing butyrate from lactate and acetate is a much lower energy-yielding process, compared to glucose [49]. Slow growth is also a known strategy to deal with starvation in other nonsporulating bacteria [58, 59].

*B. thetaiotaomicron*’s growth strategy results in a dramatic reduction in the environmental pH, which significantly inhibits its own growth. This self-inhibition resembles the hypothesis of ecological suicide suggested previously [60]. At low pH, *B. thetaiotaomicron* cells remain viable for an extended period of time when mucin beads are present. Without mucin, when glucose and pyruvate are depleted, a loss of viability is observed (Figure 2a). When the pH was restored in a spent medium assay, *B. thetaiotaomicron* metabolized an alternative carbon source and restored growth (Figure S2). Thus, the depletion of nutrients per se is probably not the cause of this viability loss. Taken together, these results suggest that similar to other bacteria [61], the metabolic activity of *B. thetaiotaomicron* requires an energy source in an acidic environment, possibly channeling its ATP from growth to maintaining its intracellular pH within a physiologically viable range. When this source of energy is no longer available, cells quickly lose viability and can no longer exploit new nutrients. Although the two species compete for glucose and pyruvate, and thus initially sustain a negative interaction, the growth of *R. intestinalis* results in a smaller decrease in the environmental pH than the growth of *B. thetaiotaomicron*, leading to an indirect positive effect on *B. thetaiotaomicron* at later stages (Figure 5).

*B. thetaiotaomicron* and *R. intestinalis* respond differently to mucin, depending also on the presence or absence of glucose. In the absence of glucose, most of the mucin sugars tested boost *B. thetaiotaomicron*’s growth, except for Neu5Ac. Analysis of its genome reveals the absence of a Neu5Ac lyase (*nanL*) gene. *B. thetaiotaomicron* is unable to catabolize Neu5Ac and did not grow on it in a minimal medium [62, 63]. As mucin beads and the tested mucin sugars did not significantly affect *B. thetaiotaomicron*’s growth during glucose fermentation (Figure S9a), we assume that glucose prevents *B. thetaiotaomicron* from simultaneously consuming other carbon sources such as mucin-derived galactose and fucose. This assumption was also necessary to reproduce *B. thetaiotaomicron*’s behavior with our model. In *E. coli*, further uptake of carbon sources is suppressed by the transcriptional regulator cAMP-Crp when nutrient uptake surpasses anabolism [53]. *B. thetaiotaomicron* uses another master regulator of carbohydrate utilization (BT_4338) particularly for mucin utilization [64], which we found to be significantly over-expressed when *B. thetaiotaomicron* consumed glucose. As *B. thetaiotaomicron* was reported to easily switch its metabolism from dietary input to mucin glycans [17], we assume that it relies on BT_4338 to cope with excessive carbon flux. In contrast, *R. intestinalis* does not grow well when glucose and mucin are supplied together (Figure 2e). Adding mucin beads also significantly up-regulated the gene of HTH-type transcriptional repressor GlcR (RIL182_03930), which indicated the repression of glucose catabolism by mucin sugars. Combined with slower growth on mucin sugars, this may explain the negative effects of mucin sugars on its growth on glucose. This is also supported by its significantly increased abundance in the mucin co-culture (Figure 5), which likely resulted from a decreased concentration of mucin sugars such as mannose due to uptake by *B. thetaiotaomicron*. However, a mechanism of *R. intestinalis*’ carbon uptake strategy has to be explored further.

Glycan-mediated attachment was reported as an important strategy for gut bacteria to access nutrients [8]. Since both *B. thetaiotaomicron* and *R. intestinalis* attached to the mucin beads, the physical proximity could influence their interactions on the beads. The beads were also not saturated by cells, illustrated by a roughly linear relationship between density on beads and in the liquid of co-culture (Figures S12 and S14). In addition, the observed fast attachment and detachment implies that the dynamics between attached and free-swimming cells may be tightly coupled. Spatial transcriptomics at a single-cell resolution may further reveal the interaction mechanisms of human gut bacteria in a spatial context [65].

In the presence of *B. thetaiotaomicron*, *R. intestinalis* exhibited additional density peaks at 38 h without mucin and at 120 h with mucin, which were not predicted by our model. In most of the cases, the classification technique we used showed high accuracy in distinguishing *B. thetaiotaomicron* and *R. intestinalis* on a basis of their optical characteristics in flow cytometry, and the method accuracy was confirmed with the data of a GFP labelled *B. thetaiotaomicron* (Figure S15). However, when the cells are losing viability (Figure 2c, 24-38 h), the similarity between live and inviable cells causes difficulties in species separation. Thus, the transition from the first peak of viable *R. intestinalis* cells to the second in the WC co-culture may be smoother than observed. There are also alternative explanations for the peak. Since *B. thetaiotaomicron* can produce a higher concentration of lactate than *R. intestinalis* in monoculture, a transiently sufficient amount of lactate in the presence of *B. thetaiotaomicron* may reverse the flow from lactate to pyruvate in *R. intestinalis* so that lactate is consumed instead of produced. In addition, under the stress of an acidic pH *B. thetaiotaomicron* may release toxins that contribute to the loss of viability of *R. intestinalis* [66]. Some viable cells of *R. intestinalis* may also grow by necrophagy on accumulated dead cells [67].

We found that interaction strength significantly varies in value and sign along the growth curve, both with and without mucin (Figure 5), due to the change in pH and nutrient levels caused by bacterial growth. Such an interplay between microbial growth and the environment is not only limited to the human gut, but also commonly described for other natural or engineered ecosystems, such as soil [68] or anaerobic digesters [69]. Understanding it requires new experimental and analytical approaches that track ecosystems in time and reveal how and when interactions change with the environment. However, widely used population models such as the generalized Lotka-Volterra assume interaction strengths to be constant. Thus, the generalized Lotka-Volterra is only applicable in systems where environmental conditions (including pH and nutrient levels) are kept constant, i.e., chemostats. The intestine is not well described by a chemostat, since nutrient levels, moisture and pH fluctuate with the daily intake of meals and the circadian rhythm [70, 71]. As confirmed by *in situ* measurements with capsules, the pH is also known to be variable across colon segments and across colons [72]. To cope with this variability, models can either account for changing interaction strengths as done here, which requires knowledge of interaction mechanisms and parameter values, or average over interaction strengths encountered in typical scenarios in the environment of interest. Such an ensemble approach may better represent the interaction between two organisms than the usual “snapshot” strategy of quantifying interactions in one particular condition but may not lend itself to predictive modelling. Thus, changing interaction strengths are a challenge for predictive models of the human gut microbiota and other ecosystems that sustain dynamic environmental conditions.

While our experiments were designed to mimic some aspects of the human colon, they neglected others. Not only is the chemical environment in the colon more complex than WC medium, but also the commercially available pig gastric mucins and subsequent treatment (e.g., autoclavation) alters the composition and structural properties compared to the colonic mucins, which could potentially alter the attachment and further utilization abilities of gut bacteria. In addition, nutrient availability and pH may vary less in the colon than in our experiments since sorption across the epithelium reduces the accumulation of fermentation products.

A deeper understanding of alternative metabolic strategies and changing species interactions in gut microbiota is necessary to rationally design next-generation probiotics and to predict the outcome of fecal microbiota transplantation. By deciphering the interaction mechanisms between two common gut bacteria across changing conditions, we have taken another step towards this goal.

## Data Availability

Raw amplicon sequencing data and RNA-seq data have been deposited to the Sequence Read Archive database under accession numbers of SAMN32182740-827 and SAMN32321133-55, respectively (https://www.ncbi.nlm.nih.gov/bioproject/PRJNA911421/ and https://www.ncbi.nlm.nih.gov/bioproject/PRJNA914119/). Raw flow cytometry data have been deposited to flowrepository.org, under Repository IDs of FR-FCM-Z6YM, FR-FCM-Z6YN, FR-FCM-Z6YP, FR-FCM-Z6YQ, FR-FCM-Z6YR, FR-FCM-Z6YS, FR-FCM-Z6YT, FR-FCM-Z6YU, FR-FCM-Z6YV and FR-FCM-Z6YX. The processed RNA-seq data, flow cytometry data and fermentation data are available at: https://github.com/msysbio.

## Code Availability

Flow cytometry data processing: bit.ly/3WNrslL, MATLAB file for the kinetic model and RNA-seq processing scripts: https://github.com/msysbio.

## Acknowledgments

We would like to thank Emma Hernandez-Sanabria, Veronica Lloréns-Rico and Jelle Matthijnssens for input and support concerning bacterial transcriptomics. In addition, we thank Vitor Pinheiro, Lieve Vanmellaert and Charlotte van de Velde for their help in constructing the GFP tagged *B. thetaiotaomicron* strain as well as Jeroen Raes for access to the CytoFLEX. We are also grateful to Thi Thuy Duyen Nguyen, Leen Rymenans and Geert Huys of the Raes lab for assistance in flow cytometry, 16S rRNA gene library preparation and anaerobic cultivation, as well as to Raul Yhossef Tito Tadeo for processing raw sequence reads. We also thank Tobie Martens from MULTIPHOTON of KU Leuven for his technical assistance in microscopy and the team of Genomics Core and Metabolomics Core of KU Leuven for data acquisition. This work was supported by funding from the European Research Council (ERC) under the European Union’s Horizon 2020 research and innovation program under grant agreement number 801747 (EcoBox) and by KU Leuven Internal funding for Interdisciplinary Networks (ID-N) PlasticDaphnia.

## Author Contributions

B.L., D.R.G. and K.F. designed the experiments and B.L. and D.R.G. carried them out. A.K. processed raw RNA-seq data. D.G. developed the kinetic model and performed simulations. K.S. measured metabolite concentrations. A.G. and K.B. gave advice on experimental design and interpretation. B.L., D.R.G., K.F. and D.G. analyzed the results and B.L., D.R.G. and K.F. wrote the manuscript. All authors read the manuscript.

## Supplementary Information

Supplementary file 1: supplementary figures S1-S15.

Supplementary file 2: description of the kinetic model.

Supplementary file 3: RNA expression data for *Bacteroides thetaiotaomicron*.

Supplementary file 4: RNA expression data for *Roseburia intestinalis*.

