## Supplementary file 1 for "Starvation responses impact interaction dynamics of human gut bacteria *Bacteroides thetaiotaomicron* and *Roseburia intestinalis*"

**This PDF file includes:**

Figures S1 to S15
SI References

**Other supporting materials for this manuscript include the following:**

Supplementary file 2: description of the kinetic model
Supplementary file 3: RNA expression data for *Bacteroides thetaiotaomicron*
Supplementary file 4: RNA expression data for *Roseburia intestinalis*

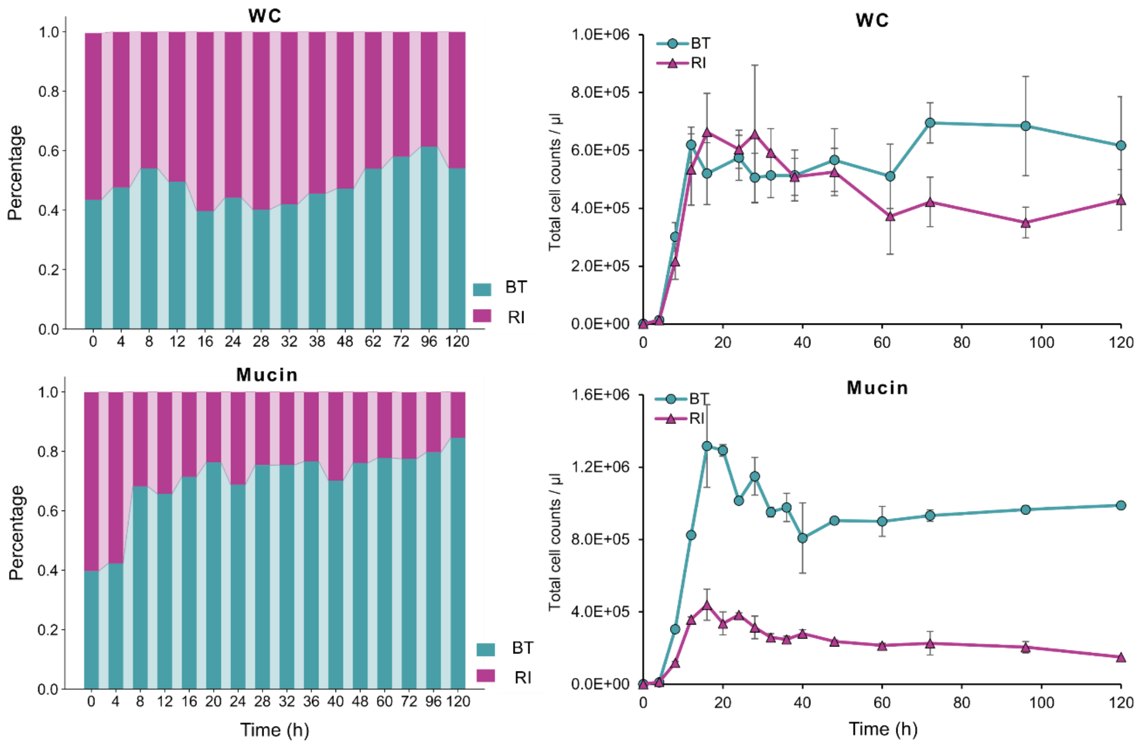


**Figure S1. Comparison of abundance profiles of *B. thetaiotaomicron* and *R. intestinalis* in co-culture with 16S rRNA gene sequencing and total cell counts of flow cytometry data.** Error bars indicate the standard deviation of four biological replicates.

BT: *Bacteroides thetaiotaomicron;* RI: *Roseburia intestinalis*; WC: Wilkins Chalgren medium.

a

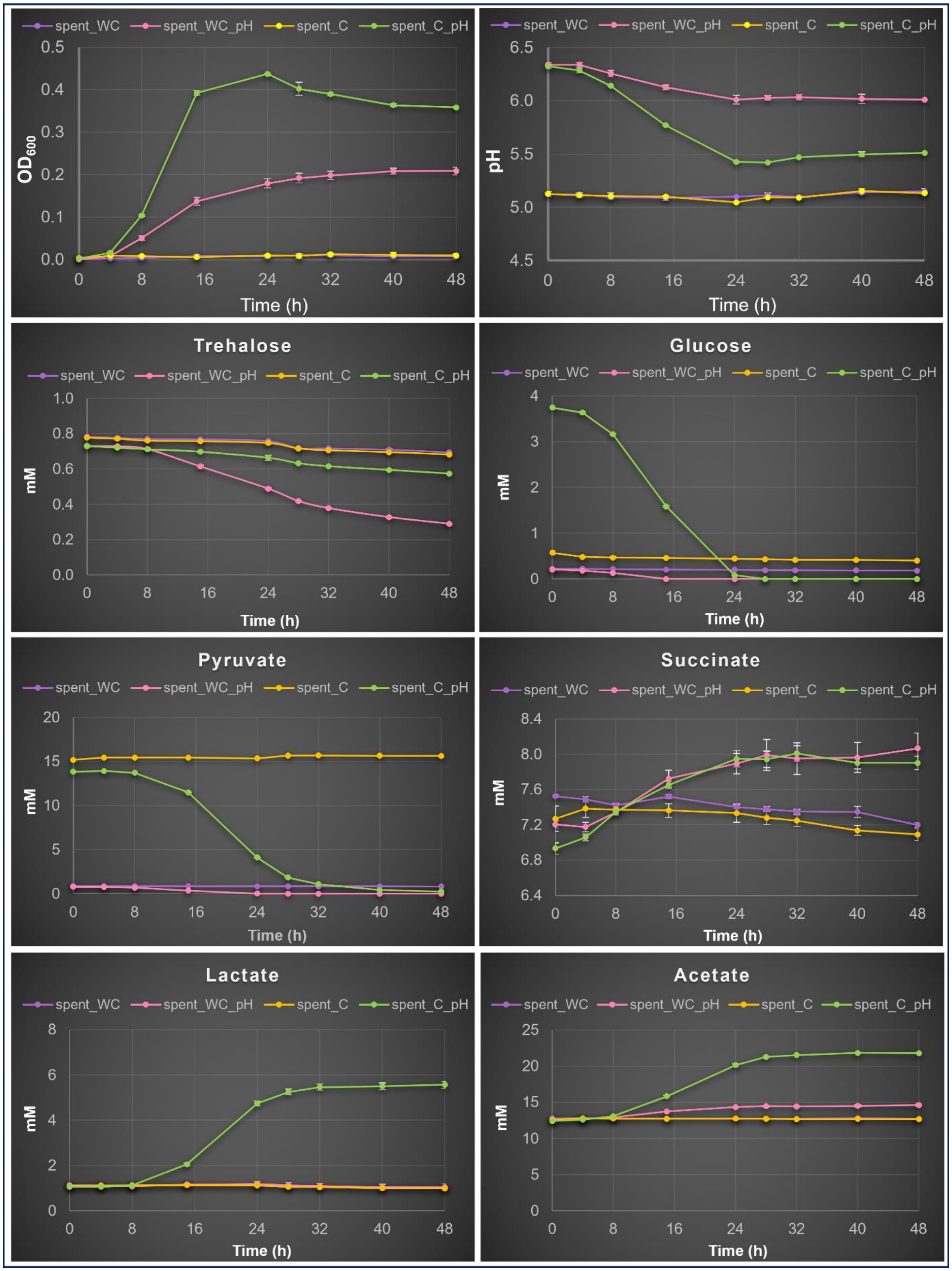

spent_WC
spent_WC_pH
spent_C
spent_C_pH
$OD_{600}$
Time (h)
pH
Trehalose
Glucose
Pyruvate
Succinate
Lactate
Acetate
mM

**b**

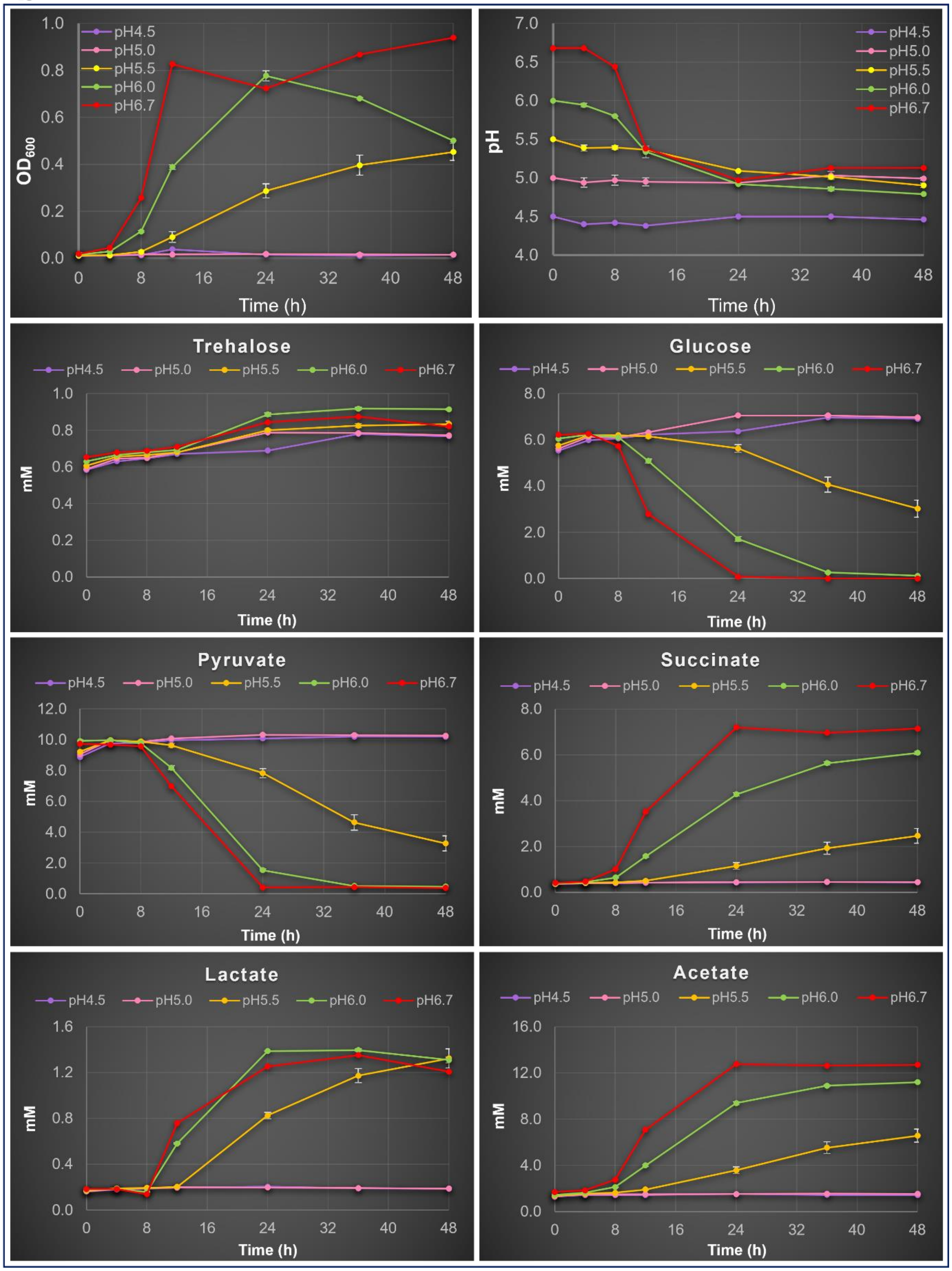


**Figure S2. Cells of *B. thetaiotaomicron* lose viability at acidic pH conditions in serum bottles. a**, *B. thetaiotaomicron* re-grown in its spent media collected at 48 h cultivation after nutrient depletion. spent_WC: the original spent medium without adjusting pH (pH: 5) and without carbon source added; spent_WC_pH: spent medium with pH adjusted to 6.5; spent_C: spent medium with glucose and pyruvate added in the same concentrations as in WC (Wilkins Chalgren medium) at pH 5; spent_C_pH: spent medium with added glucose and pyruvate plus pH adjusted to 6.5. **b**. *B. thetaiotaomicron* grown in WC media with pH values adjusted to 4.5, 5.0, 5.5, 6.0 and 6.7. Error bars indicate the standard deviation of three biological replicates.

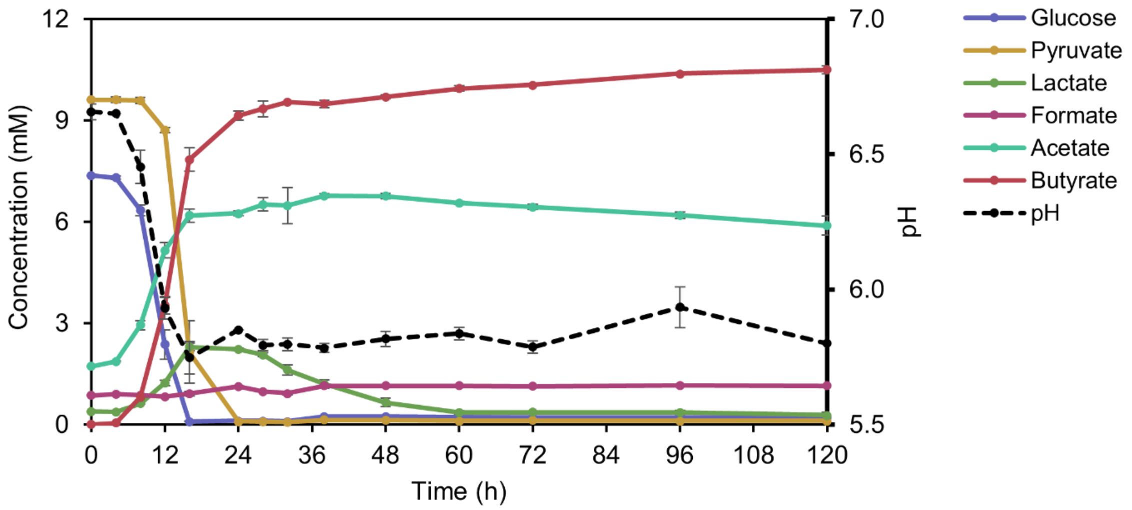


**Figure S3. Consumption of lactate in *R. intestinalis* monoculture.** After the depletion of glucose and pyruvate in WC, *R. intestinalis* enters a slow growth mode where it converts lactate and acetate to butyrate. Error bars indicate the standard deviation of three biological replicates in bottle experiment.

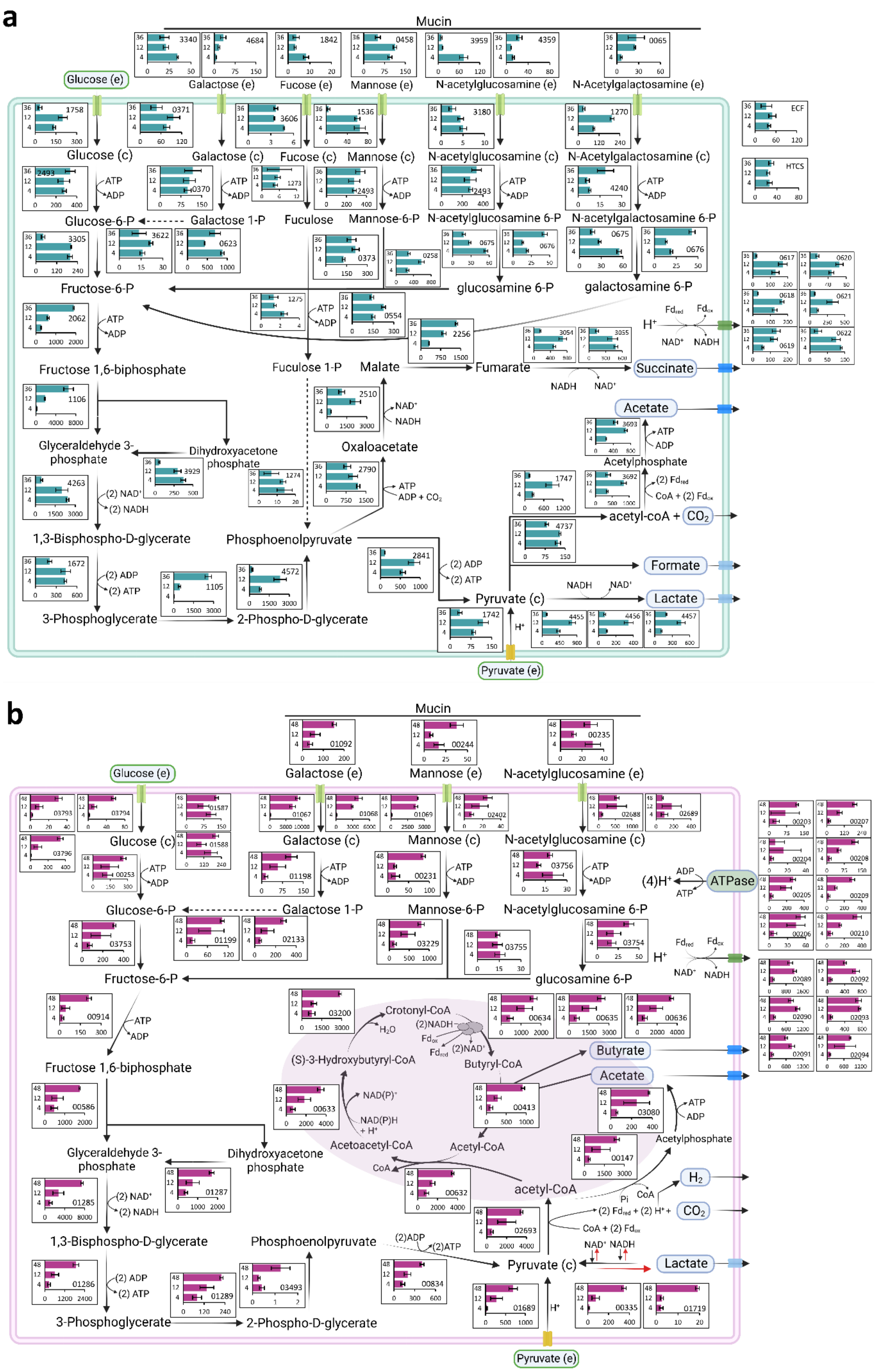
a
Mucin
Glucose (e)
Galactose (e)
Fucose (e)
Mannose (e)
N-acetylglucosamine (e)
N-Acetylgalactosamine (e)
ECF
HTCS
Glucose (c)
Galactose (c)
Fucose (c)
Mannose (c)
N-acetylglucosamine (c)
N-Acetylgalactosamine (c)
ATP
ADP
Glucose-6-P
Galactose 1-P
Fuculose
Mannose-6-P
N-acetylglucosamine 6-P
N-acetylgalactosamine 6-P
Fructose-6-P
glucosamine 6-P
galactosamine 6-P
Fructose 1,6-biphosphate
Fuculose 1-P
Malate
Fumarate
Succinate
Acetate
Glyceraldehyde 3-phosphate
Dihydroxyacetone phosphate
Oxaloacetate
Acetylphosphate
1,3-Bisphospho-D-glycerate
Phosphoenolpyruvate
acetyl-coA + CO2
Formate
Pyruvate (c)
Lactate
3-Phosphoglycerate
2-Phospho-D-glycerate
Pyruvate (e)
NAD+
NADH
(2) NAD+
(2) NADH
(2) ADP
(2) ATP
ADP + CO2
H+
b
Mucin
Glucose (e)
Galactose (e)
Mannose (e)
N-acetylglucosamine (e)
Glucose (c)
Galactose (c)
Mannose (c)
N-acetylglucosamine (c)
(4)H+
ATPase
Glucose-6-P
Galactose 1-P
Mannose-6-P
N-acetylglucosamine 6-P
Fructose-6-P
glucosamine 6-P
Crotonyl-CoA
(S)-3-Hydroxybutyryl-CoA
Butyryl-CoA
Butyrate
Acetate
Fructose 1,6-biphosphate
Acetoacetyl-CoA
Acetyl-CoA
Acetylphosphate
Glyceraldehyde 3-phosphate
Dihydroxyacetone phosphate
acetyl-CoA
H2
CO2
1,3-Bisphospho-D-glycerate
Phosphoenolpyruvate
Pyruvate (c)
Lactate
3-Phosphoglycerate
2-Phospho-D-glycerate
Pyruvate (e)

**Figure S4. Metabolic maps summarizing gene expression in *B. thetaiotaomicron* and *R. intestinalis* monocultures in WC.** Gene expression of the key metabolic routes of glucose and pyruvate fermentation, as well as mucin hydrolysis and mucin sugar utilization in *B. thetaiotaomicron* **(a)** and *R. intestinalis* **(b)** monocultures in WC without mucin beads. In each barplot of **a** and **b**, gene expression levels (x axis) were measured in Transcripts Per Million (TPM) of RNA samples at time (y axis) 4 h, 12 h and 36 h for *B. thetaiotaomicron*, and at time 4 h, 12h and 48 h for *R. intestinalis*. Numbers in each barplot stand for the Gene ID in genomes of *B. thetaiotaomicron* and *R. intestinalis*. The solid and dashed lines represent single- and multi-enzyme reactions between the two indicated molecules, respectively. Error bars indicate the standard deviation of three biological replicates.

ECF: extracytoplasmic function sigma/anti-sigma pair; HTCS: hybrid two-component system.

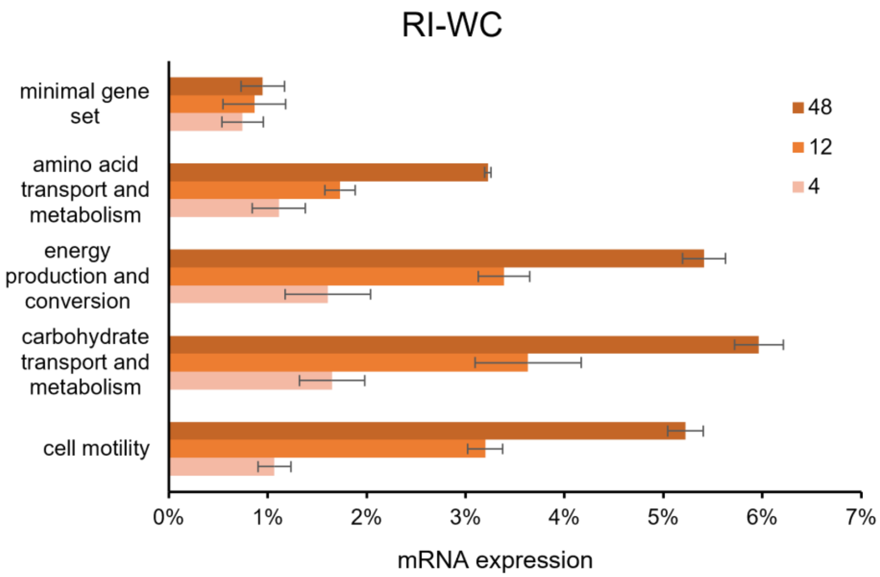


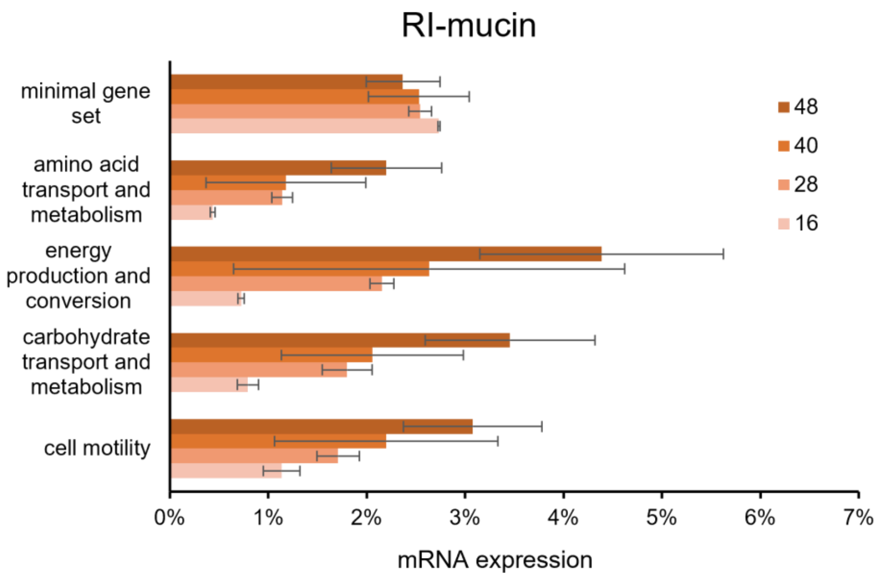


**Figure S5. *R. intestinalis* keeps expressing non-essential genes after glucose depletion.** Several major gene categories at 28 h, 40 h and 48 h are significantly ($p < 0.001$ in all cases; student's t test) up-regulated compared to at 4 h. Expression values of minimal gene sets [1] did not change significantly over time ($p > 0.05$; student's t test). mRNA abundances determined by RNA-seq are represented as percent of total mRNA. Genes are categorized using the EggNOG classification. The minimal gene set includes well-conserved housekeeping genes for basic metabolism and macromolecular synthesis, and was computed using the MicroScope [2] platform. Numbers in the legend stand for the sampling time in hours. Error bars indicate the standard deviation of three biological replicates. RI: *Roseburia intestinalis*; WC: Wilkins Chalgren medium.

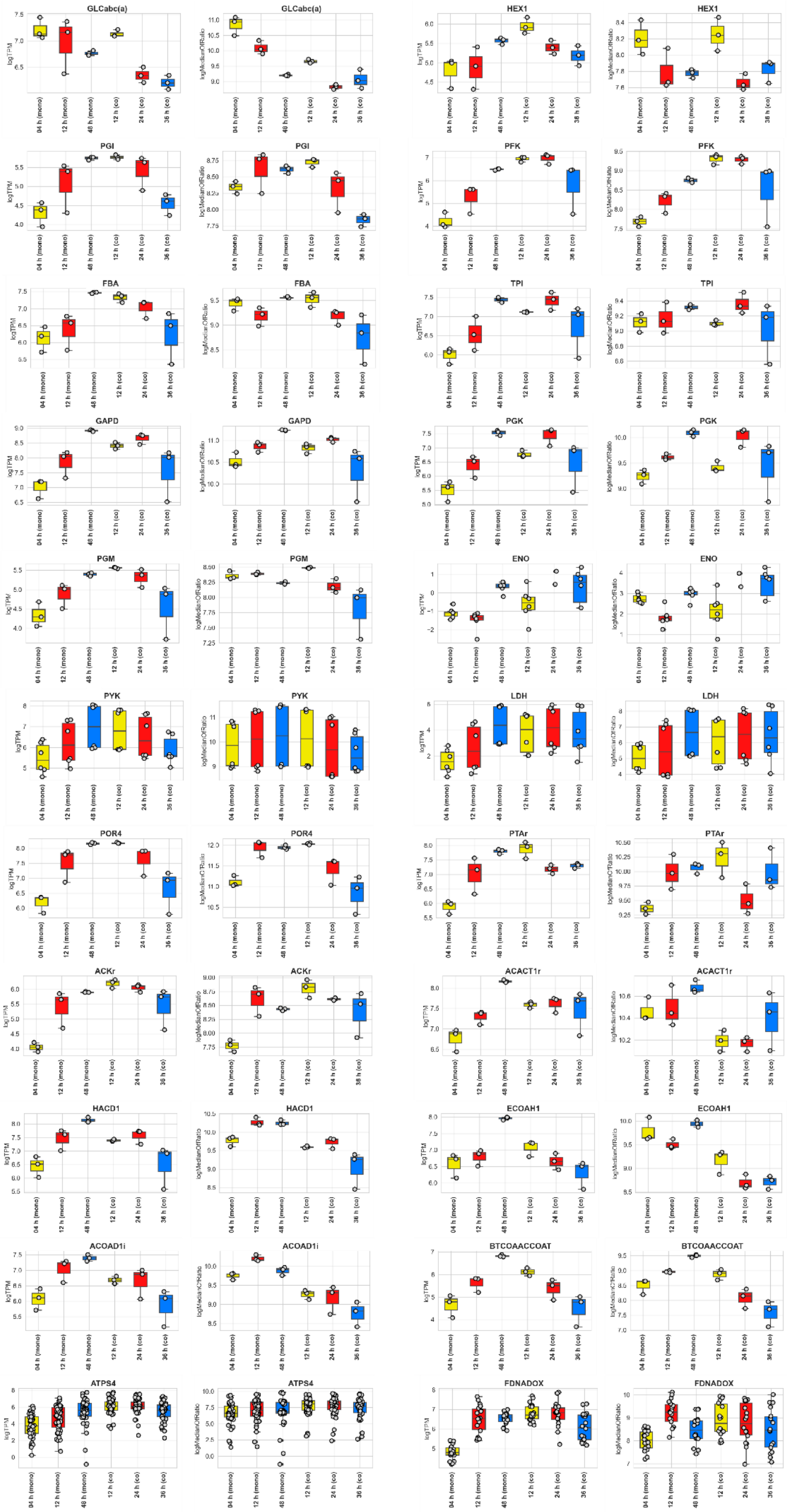
GLCabc(a)
HEX1
PGI
PFK
FBA
TPI
GAPD
PGK
PGM
ENO
PYK
LDH
POR4
PTAr
ACKr
ACACT1r
HACD1
ECOAH1
ACOAD1i
BTCOAACCOAT
ATPS4
FDNADOX
logTPM
logMedianOfRatio
04 h (mono)
12 h (mono)
48 h (mono)
12 h (co)
24 h (co)
36 h (co)

**Figure S6. Gene expression of *R. intestinalis* shows consistent results between two normalization methods.** We compared (logarithmic) gene expression values of *R. intestinalis* over time quantified as Transcripts Per Million versus DESeq2's median of ratios for selected enzymes. The name of each chart stands for the reaction ID of glycolysis. Details about the gene expression of each reaction can be found in the Supplementary file 4 (*R. intestinalis*).

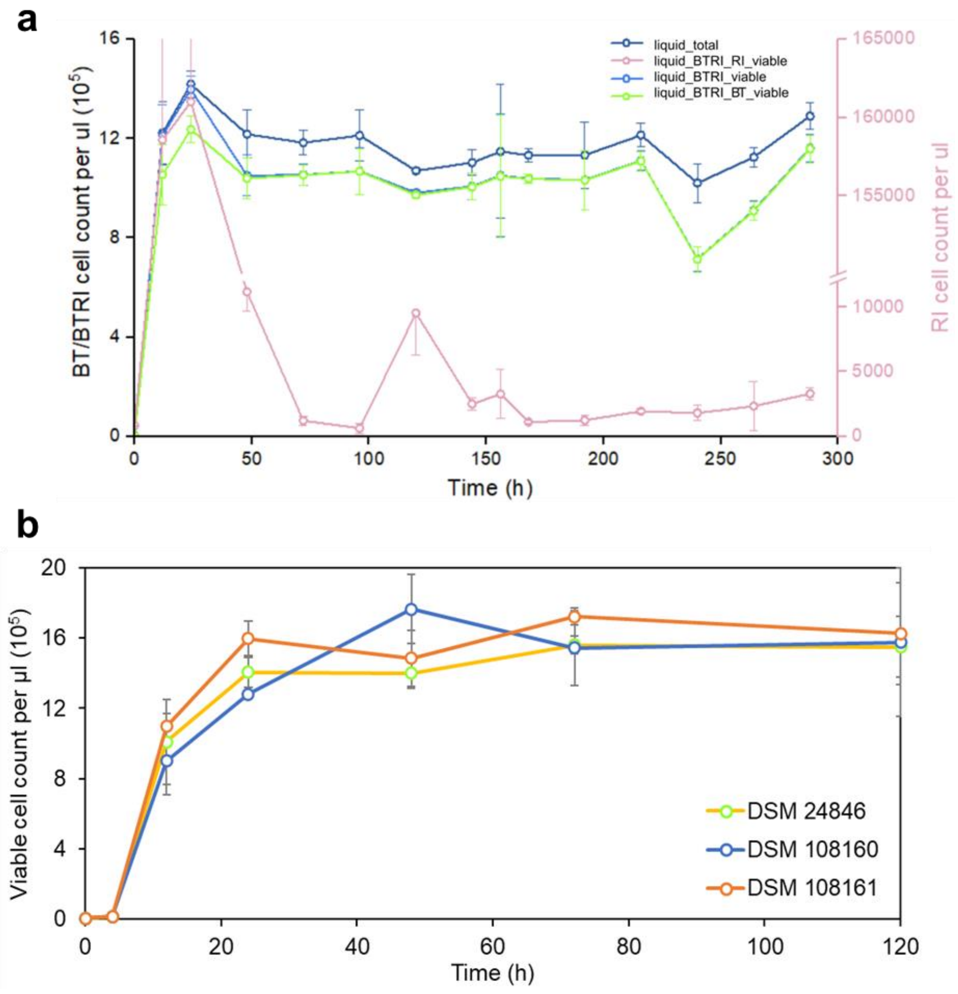


**Figure S7. *B. thetaiotaomicron* survives long term in the presence of mucin beads in bottles. a**) The co-culture of *B. thetaiotaomicron* and *R. intestinalis* was followed for 288 h with samples taken every 24 h starting from 120 h after inoculation. Data shown before 120 h were from the experiment of co-culture in WC plus mucin beads. **b**) Other three non-type strains of *B. thetaiotaomicron* also exhibited long-term growth after depleting glucose in WC supplemented with mucin beads. Error bars indicate the standard deviation of three biological replicates. BT: *Bacteroides thetaiotaomicron;* RI: *Roseburia intestinalis.*

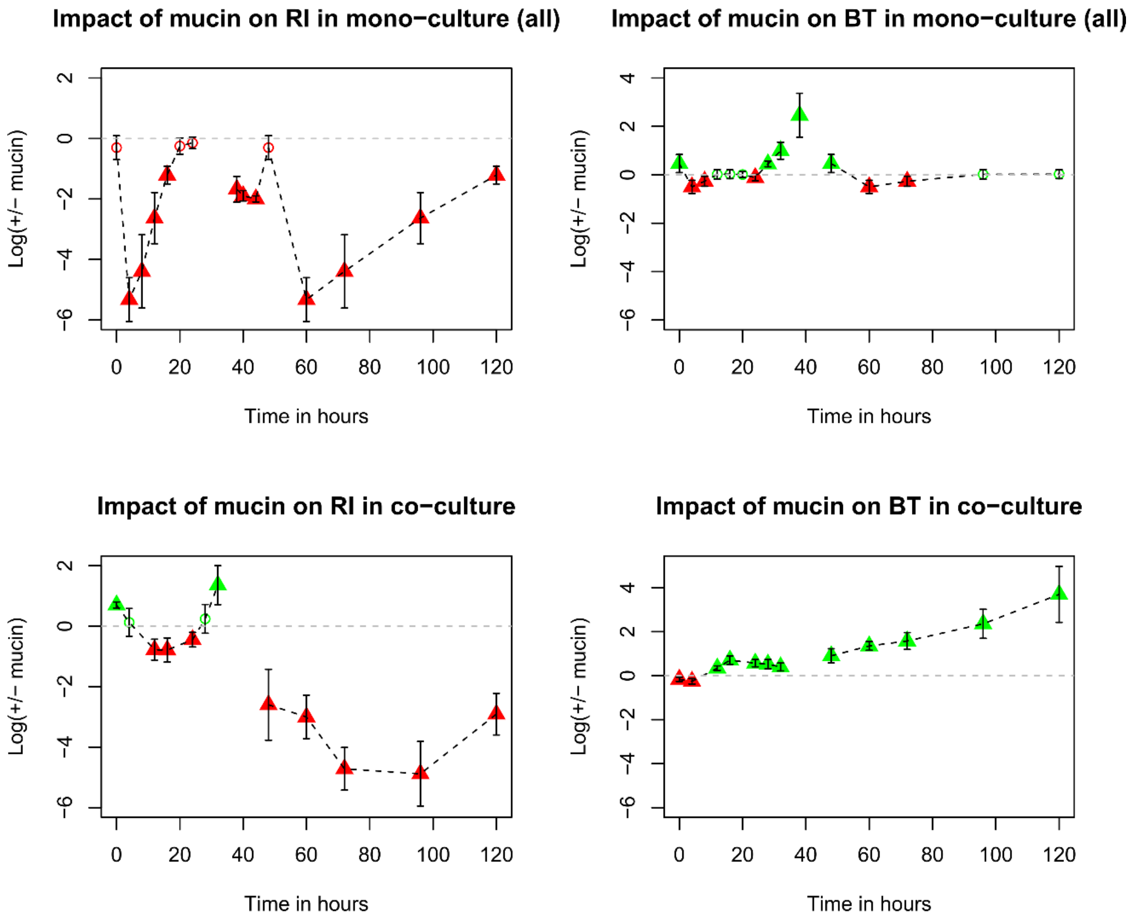


**Figure S8. Impact of mucin on growth of *B. thetaiotaomicron* and *R. intestinalis*.** We used viable cell counts in WC with mucin beads and WC without mucin beads over time to compute the impact of mucin on abundances of *R. intestinalis* and *B. thetaiotaomicron*, as the log ratio of viable cell counts in liquid in WC with mucin (+) versus WC without mucin (-). Triangle: significant Wilcoxon *p* value after Benjamini-Hochberg multiple testing correction. Mean and standard deviation of viable cell counts in liquid are computed on all possible mucin (+) versus mucin (-) pairs across replicates per time point. We assessed the impact of mucin for both mucin-free monoculture experiments together (all). BT: *Bacteroides thetaiotaomicron;* RI: *Roseburia intestinalis*; WC: Wilkins Chalgren medium.

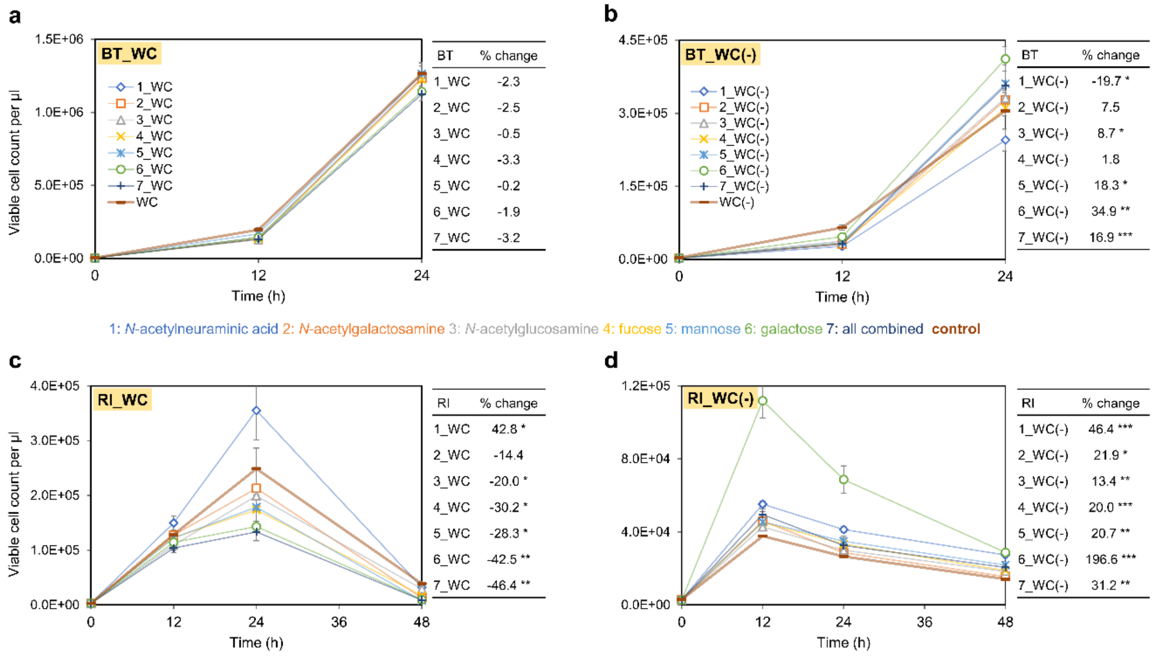


**Figure S9. Mucin sugars impact the growth of *B. thetaiotaomicron* and *R. intestinalis* differently in the presence or absence of glucose.** WC and WC(-) represent the standard WC medium and WC medium without carbon sources glucose and pyruvate. Each medium was split into eight aliquots, each of which was supplemented with either one of the listed mucin sugars or a mixture of all six sugars together or without any sugar added as the control. Error bars indicate the standard deviation of three biological replicates in 24-well plates. Quantitative information of positive and negative effects of mucin sugars are shown in tables (% change compared to the control). Asterisks: when cell density in the mucin sugar group is significantly different than in the control (two-tailed t-test; * $p < 0.05$, ** $p < 0.01$, *** $p < 0.001$).

BT: *Bacteroides thetaiotaomicron;* RI: *Roseburia intestinalis*; WC: Wilkins Chalgren medium.

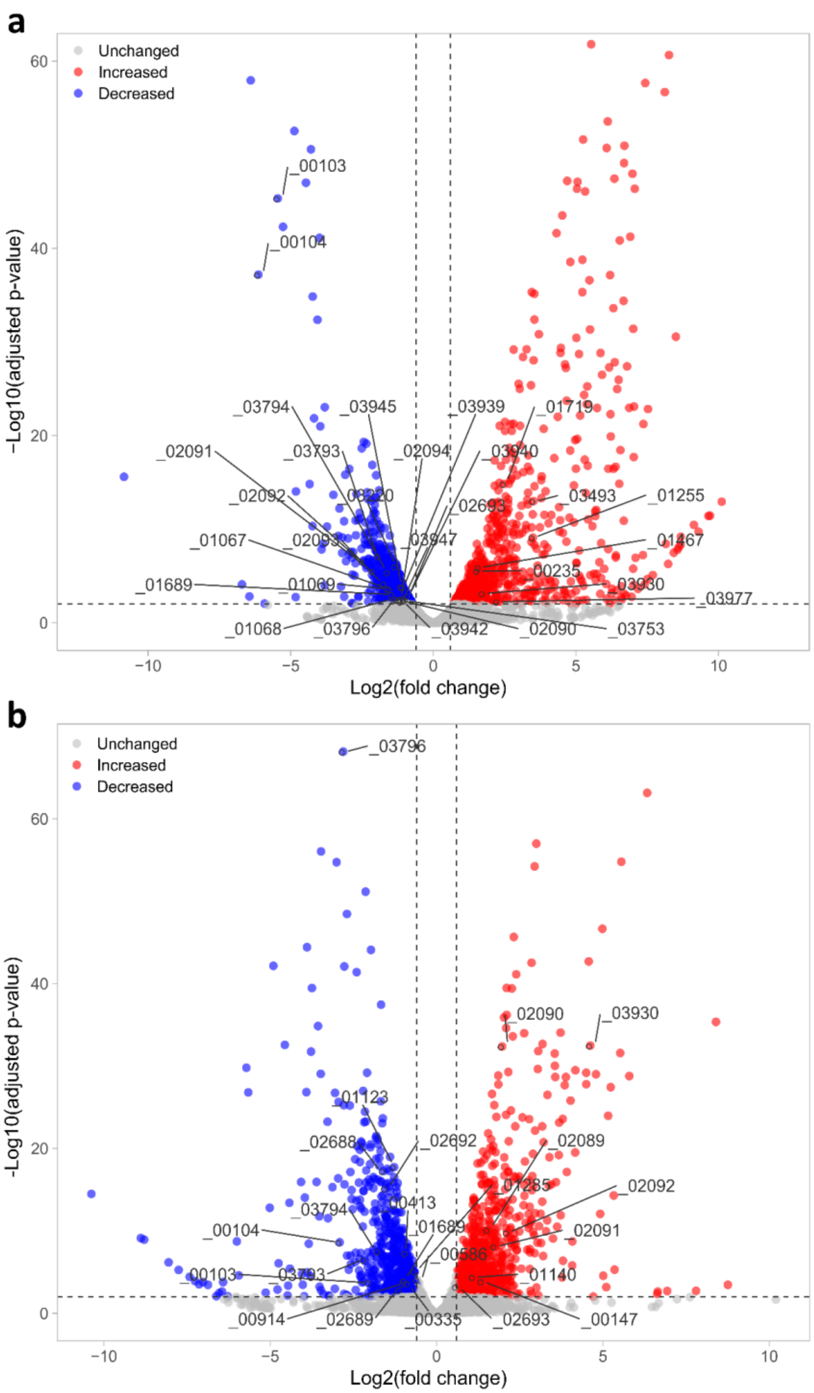


**Figure S10. Differentially expressed genes in *R. intestinalis* monoculture without and with mucin beads.** We calculated the differential expression of genes between *R. intestinalis* grown in WC monoculture and WC monoculture with mucin beads at high glucose/pyruvate levels (**a**) and low glucose/pyruvate levels (**b**). Numbers represent the Gene ID in the genome of *R. intestinalis*. We used a strict cutoff of at least 1.5-fold and a Benjamini-Hochberg adjusted *p* value less than or equal to 0.01 for differential expression calculated by DESeq2.

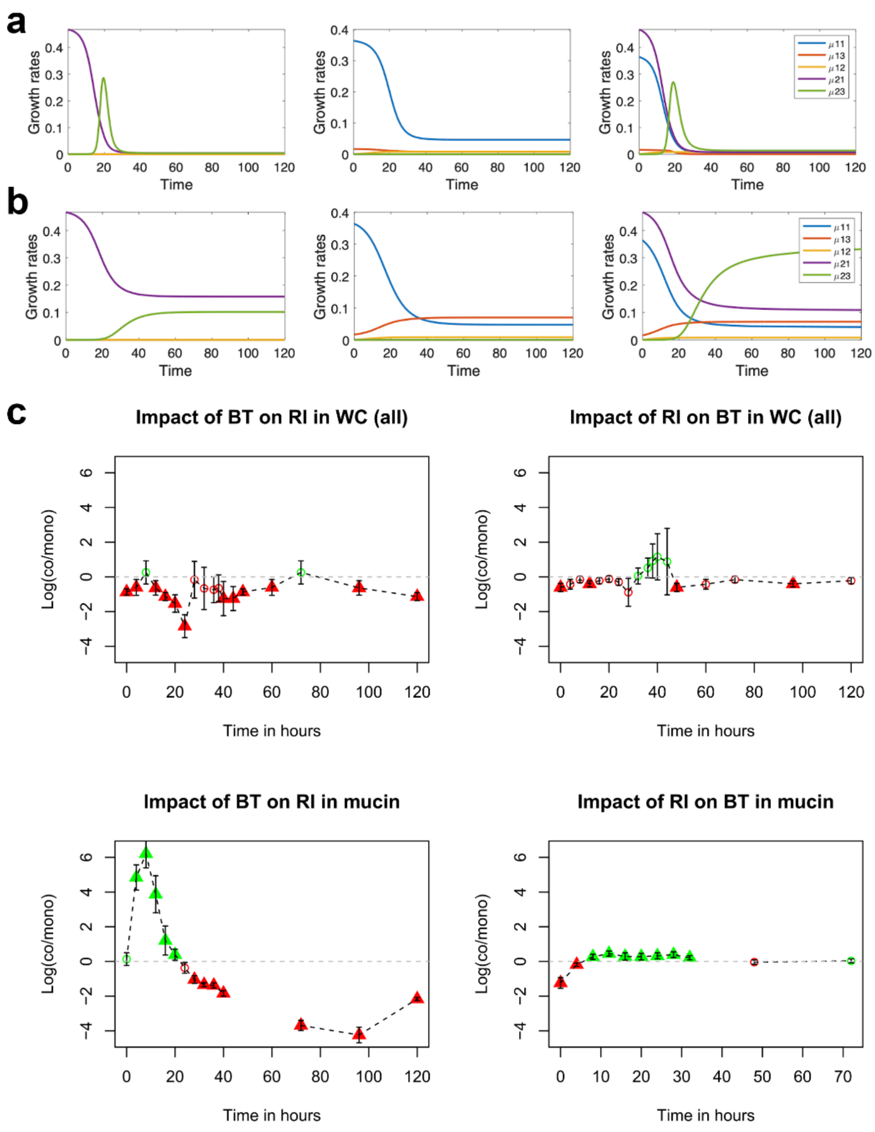


**Figure S11. Interaction strength and sign changes over time for *B. thetaiotaomicron* and *R. intestinalis* with or without mucin. a**) Growth rates of *B. thetaiotaomicron* (left), *R. intestinalis* (middle) and their co-culture (right) in WC without mucin predicted with the kinetic model. **b**) Growth rates of *B. thetaiotaomicron* (left), *R. intestinalis* (middle) and their co-culture (right) in WC with mucin predicted with the kinetic model. **c**) Interaction strength and sign of *B. thetaiotaomicron* and *R. intestinalis* with and without mucin. Interaction strength is sometimes quantified as the ratio of growth rates in exponential phase (especially in the context of metabolic modelling) [3–5], which is not appropriate here because of the slow growth mode. We therefore computed interaction strength, i.e., the impact of one organism on the other's abundance, as the log ratio of viable cell counts in liquid in co- versus monocultures, which is also common in the literature [6, 7]. µ11: growth rate of *R. intestinalis* on glucose, µ12: growth rate of *R. intestinalis* on lactate/acetate, µ13: growth rate of *R. intestinalis* on mucin sugars, µ21: growth rate of *B.*

*thetaiotaomicron* on glucose, µ23: growth rate of *B. thetaiotaomicron* on mucin sugars. Triangle: significant Wilcoxon *p* value after Benjamini-Hochberg multiple testing correction. Mean and standard deviation of viable cell counts in liquid are computed on all possible co- versus monoculture pairs across replicates per time point. For *B. thetaiotaomicron* and *R. intestinalis* without mucin, we calculated interaction strength from data of both monoculture experiments together (all). Predicted growth rates indicate mechanisms behind changing interaction strengths. For instance, *B. thetaiotaomicron* grows better on the residual sugars in the presence of *R. intestinalis*. It is of note that the residual sugars including some mucin sugars are present in low amounts also in the absence of mucin beads. BT: *Bacteroides thetaiotaomicron;* RI: *Roseburia intestinalis*; WC: Wilkins Chalgren medium.

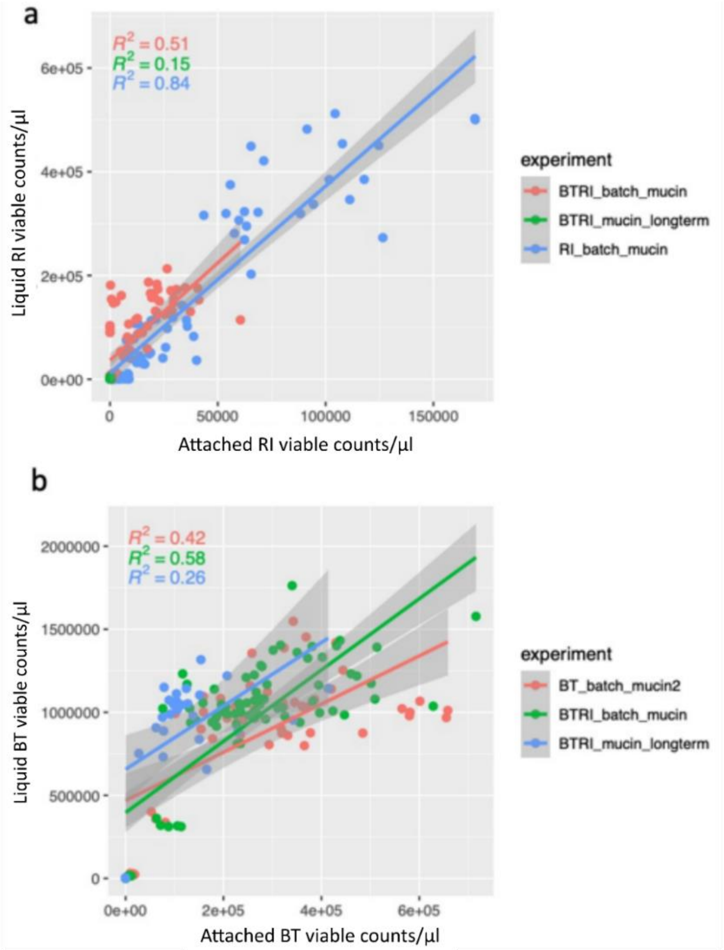


**Figure S12. Counts of viable attached and planktonic cells scale roughly linearly (Pearson correlation). a**) relationship between viable cell counts of *R. intestinalis* in liquid and attached to the mucin beads. **b**) relationship between viable cell counts of *B. thetaiotaomicron* in liquid and attached to the mucin beads.

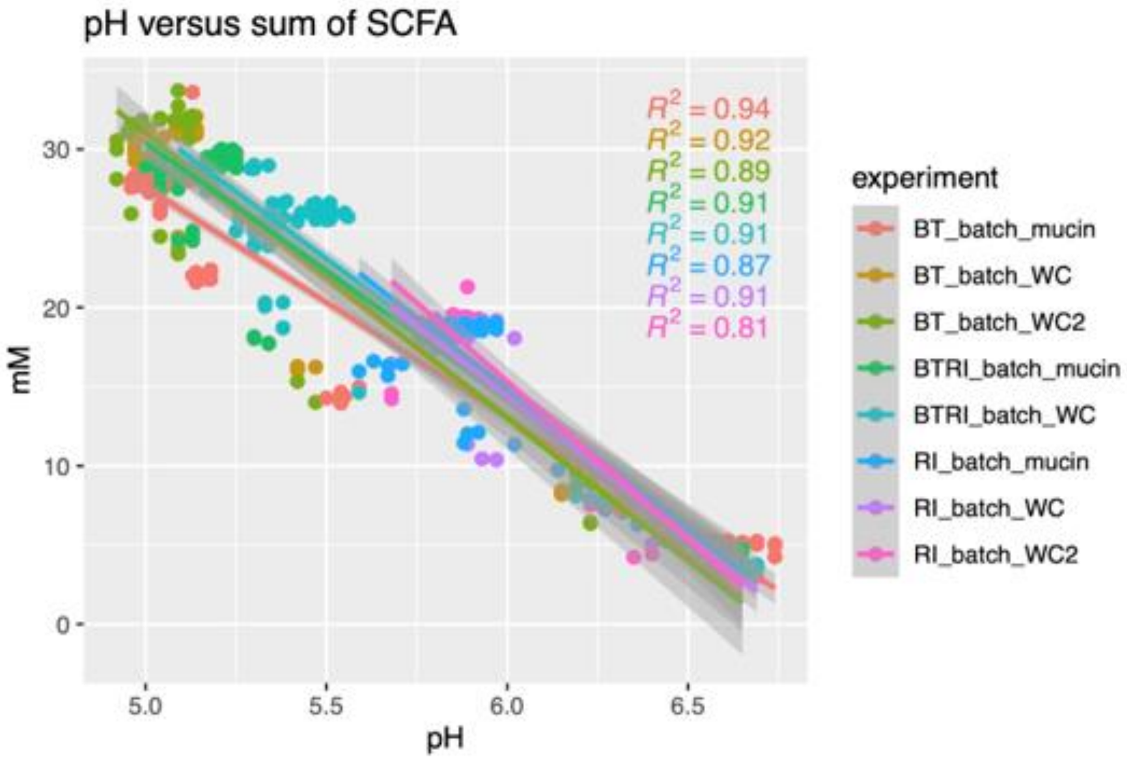


**Figure S13. pH scales linearly (Pearson correlation) with the sum of the concentrations of short-chain fatty acids (SCFA).** BT: *Bacteroides thetaiotaomicron;* RI: *Roseburia intestinalis*; WC: Wilkins Chalgren medium.

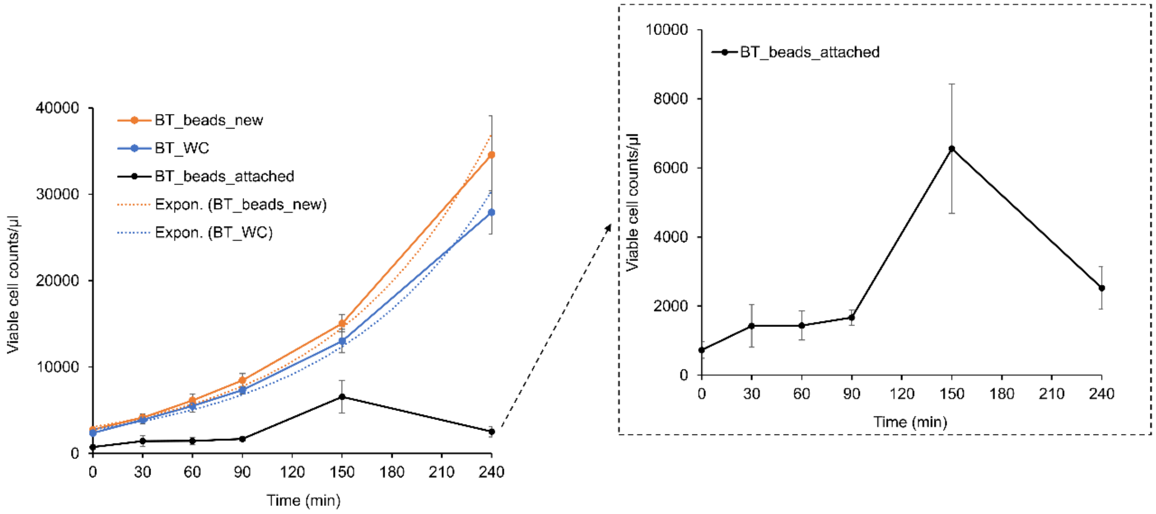


**Figure S14. Attached *B. thetaiotaomicron* cells rapidly detach from mucin beads.** After growing *B. thetaiotaomicron* in WC for 24 h, we collected liquid fermentation broth that contains mucin beads with attached *B. thetaiotaomicron* cells and used PBS for washing away the residual cells in liquid. Later mucin beads with attached *B. thetaiotaomicron* cells were added into fresh WC medium (in black). The right figure zooms in which the attached cells detached from mucin beads to the WC medium. For comparison, we included *B. thetaiotaomicron* grown in fresh WC medium (in blue) and in WC medium supplemented with sterile mucin beads (in orange). Dashed trendlines are shown in exponential scale. Error bars indicate the standard deviation of four biological replicates. BT: *Bacteroides thetaiotaomicron*; WC: Wilkins Chalgren medium.

a

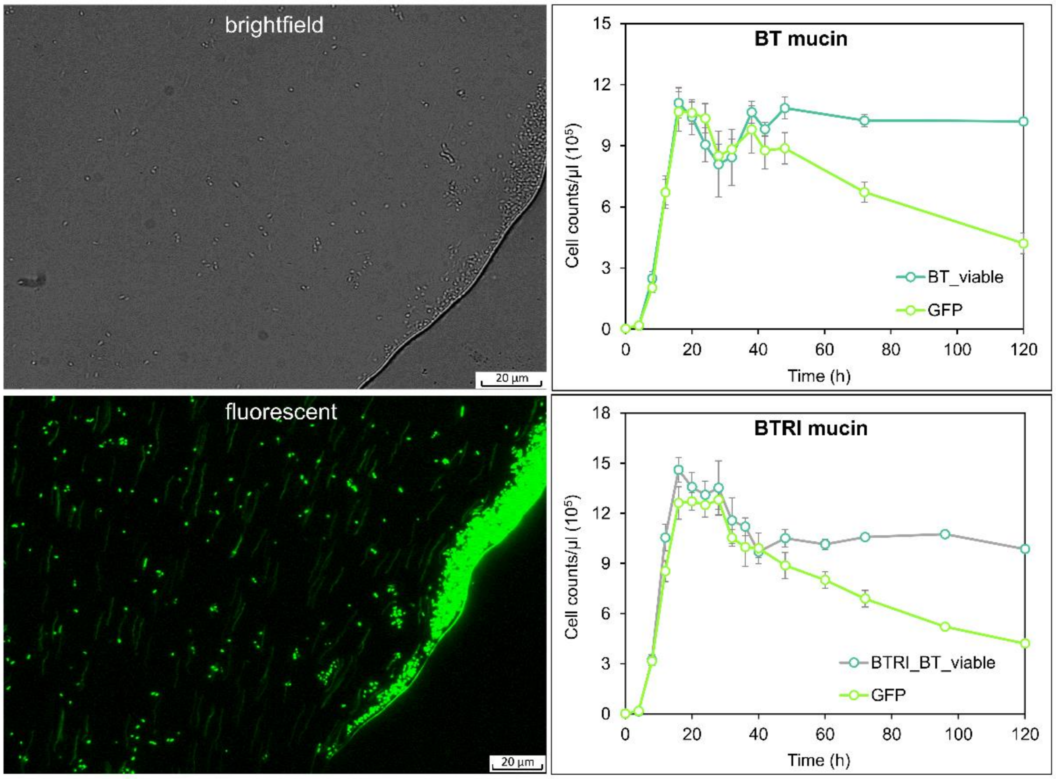
brightfield
20 µm
fluorescent
20 µm
BT mucin
Cell counts/µl ($10^5$)
Time (h)
BT_viable
GFP
BTRI mucin
Cell counts/µl ($10^5$)
Time (h)
BTRI_BT_viable
GFP

b

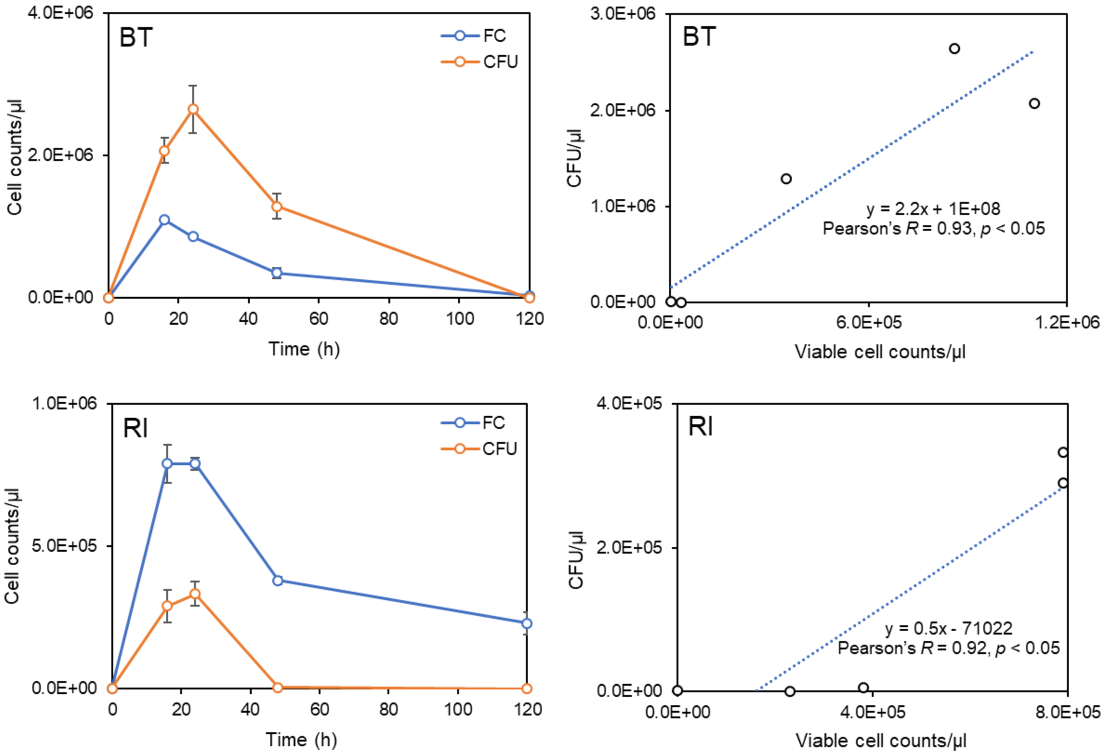
BT
FC
CFU
Cell counts/µl
Time (h)
BT
CFU/µl
Viable cell counts/µl
y = 2.2x + 1E+08
Pearson's R = 0.93, p < 0.05
RI
FC
CFU
Cell counts/µl
Time (h)
RI
CFU/µl
Viable cell counts/µl
y = 0.5x - 71022
Pearson's R = 0.92, p < 0.05

**Figure S15. Cell counts of GFP labelled *B. thetaiotaomicron* and colony-forming units confirm method accuracy of flow cytometry with UMAP. a**) GFP labelled *B. thetaiotaomicron* can stably fluoresce when grown anaerobically with subsequent exposure to oxygen (left panel); In the first 40 hours, viable cell counts of *B. thetaiotaomicron* in monoculture and co-culture both show no significant ($p$ > 0.05; Student's t test) differences compared to the cell counts of GFP labelled *B. thetaiotaomicron*, numbers of which were obtained from the FITC channel of a CytoFLEX S flow cytometer (right panel). **b**) Comparison between viable cell counts from flow cytometry and colony-forming units over time for both *B. thetaiotaomicron* and *R. intestinalis*. On the right panel, the trendlines show a linear correlation between viable cell counts and colony-forming units, with the strength and significance of Pearson's correlation displayed next to them. Error bars indicate the standard deviation of three biological replicates. BT: *Bacteroides thetaiotaomicron;* RI: *Roseburia intestinalis*; CFU: colony-forming unit.
