## Supplementary file 2 for "Starvation responses impact interaction dynamics of human gut bacteria *Bacteroides thetaiotaomicron* and *Roseburia intestinalis*"

### Modeling the dynamics of the RI/BT co-culture

#### Assumptions

In order to reproduce the dynamics of the bacterial abundances and of the nutrient concentration, we build a tentative ODE-based model. This model is based on the following assumptions (Fig. 4a):

- For each strain, we distinguish growing and slow growth modes. This latter state is characterized by low growth/death rates. Switches to slow growth states are induced by the level of available nutrients.
- A constant fraction of each strain is attached to the mucin beads. Liquid and attached bacteria grow at the same rate on a given sugar source but only the attached bacteria can degrade mucin.
- The chemical compounds considered in the model are glucose/pyruvate (lumped into a single variable), acetate/lactate (lumped into a single variable), and mucin sugars (mannose, galactose, etc) as well as other residual sugars (such as trehalose, present in the WC medium), lumped into a single variable and referred to as "mucin sugars".
- Initially, glucose is present in the growth medium, but no acetate/lactate. We also assume that a low concentration of mucin sugars is also present in the medium, even in absence of mucin.
- Both RI and BT grow on glucose and on mucin sugars. We further assume that mucin sugars inhibit growth of RI on glucose (experimental observation). Glucose inhibits growth of BT on mucin sugars (hypothesis made to reproduce the 2-peak growth of BT in monoculture).
- At high glucose concentration, both RI and BT produce acetate/lactate when growing on glucose. At low glucose/mucin sugars concentration, RI switches to the slow growth state, consuming acetate/lactate at a low rate.
- Both strains degrade mucin to produce mucin sugars, usable for growth.
- Butyrate is produced by RI when growing (regardless the nutrient source). Note that butyrate is produced but not consumed by the bacteria and thus does not impact their dynamics.
- Low pH inhibits growth of RI and of BT. It has a stronger (negative) impact on the growth of BT (i.e. BT is more sensitive to low pH).
- Changes in pH result from changes in acetate/lactate.

#### Variables

The variables as well as their initial values are listed in Table S1.

| Variable | Description | Initial value |
|---|---|---|
| $X_1$ | RI (*Roseburia intestinalis*) | 0.01 |
| $X_2$ | BT (*Bacteroides thetaiotaomicron*) | 0.01 |
| $\hat{X}_1$ | RI in slow growth state | 0 |
| $\hat{X}_2$ | BT in slow growth state | 0 |
| $\bar{X}_1$ | attached RI (fraction: $\bar{X}_1 = f_{X1} * (S_0 > 0) * X_1$) | - |
| $\bar{X}_2$ | attached BT (fraction: $\bar{X}_2 = f_{X2} * (S_0 > 0) * X_2$) | - |
| $S_0$ | mucin (constant) | 0 or 1 |
| $S_1$ | glucose/pyruvate | 1 |
| $S_2$ | acetate/lactate | 0 |
| $S_3$ | mucin sugars (mannose, galactose, etc) | 0.1 |
| $S_4$ | butyrate | 0 |
| $pH$ | implicitly taken into account via the functions $\phi_i$ | - |

Table S1: Variables of the model

**Bacteria densities**

The densities of the bacteria are affected by growth (which itself depends on nutrients), by switch to slow growth mode (induced by the level of available nutrients) and by death:

$$\text{RI:} \quad \frac{dX_1}{dt} = \mu_1 X_1 - \delta_1 X_1 - \kappa_{m1} X_1 = (\mu_1 - \delta_1 - \kappa_{m1}) X_1 \tag{1}$$

$$\frac{d\hat{X}_1}{dt} = \kappa_{m1} X_1 + \mu_{m1} \hat{X}_1 - \delta_{m1} \hat{X}_1 \tag{2}$$

$$\text{BT:} \quad \frac{dX_2}{dt} = \mu_2 X_2 - \delta_2 X_2 - \kappa_{m2} X_2 = (\mu_2 - \delta_2 - \kappa_{m2}) X_2 \tag{3}$$

$$\frac{d\hat{X}_2}{dt} = \kappa_{m2} X_2 - \delta_{m2} \hat{X}_2 \tag{4}$$

where
$\mu_i$ = growth rate of species $i$ (function of the nutrients)
$\delta_i$ = death rate of species $i$ (constant)
$\kappa_{mi}$ = rate of switch to slow growth state of species $i$ (function of the nutrients)

**Growth rates**

To describe the nutrient-dependent growth, we used (additive) Monod functions, modulated by conditional and pH functions. The generic form of the equations for the growth rate of species $i$ is:

$$\mu_i = \phi_i \sum_j v_{ij} \; f_{ij}(S_k) \; \frac{S_j}{K_{ij} + S_j} \tag{5}$$

where $S_j$ = nutrient $j$, $v_{ij}$ = maximum growth rate on compound $j$, $K_{ij}$ = Monod constant associated to growth on $S_j$, $f_{ij}(S_k)$ is a (facultative) conditional function of nutrient $k$ (e.g. diauxic shift), and $\phi_i$ describes phenomenologically the negative effect of pH (function of the acids).

More specifically, for RI and BT, the growth functions read as follows:

$$\mu_1 = \phi_1 \Big( \underbrace{v_{11} f_{inhib1} \frac{S_1}{K_{11}+S_1}}_{\substack{\mu_{11} \\ \text{growth on glucose} \\ \text{inhibited by mucin sugars}}} + \underbrace{v_{13} \frac{S_3}{K_{13}+S_3}}_{\substack{\mu_{13} \\ \text{growth on} \\ \text{mucin sugars}}} \Big) \tag{6}$$

$$\mu_{m1} = \phi_1 \underbrace{v_{12} \frac{S_2}{K_{12}+S_2}}_{\substack{\mu_{12} \\ \text{slow growth on acetate/lactate} \\ \text{at low glucose/mucin sugars}}} \tag{7}$$

$$\mu_2 = \phi_2 \Big( \underbrace{v_{21} \frac{S_1}{K_{21}+S_1}}_{\substack{\mu_{21} \\ \text{growth on glucose}}} + \underbrace{v_{23}\ f_{inhib2} \frac{S_3}{K_{23}+S_3}}_{\substack{\mu_{23} \\ \text{growth on other mucin sugars} \\ \text{inhibited by glucose}}} \Big) \tag{8}$$

Regulatory functions:

$$f_{inhib1} = \frac{K_{i11}}{K_{i11}+S_3} \quad \text{(inhibition of RI growth by mucin sugars when growing on glucose)} \tag{9}$$

$$f_{inhib2} = \frac{K_{i13}^n}{K_{i13}^n+S_1^n} \quad \text{(inhibition of BT growth by glucose when growing on mucin sugars)} \tag{10}$$

Rates of switches to slow growth mode:

$$\kappa_{m1} = v_{m1} \frac{K_{i1}^h}{K_{i1}^n+S_1^h} \frac{K_{i3}^h}{K_{i3}^n+S_3^h} \quad \text{(switch of RI to slow growth mode at low glucose/mucin sugar)} \tag{11}$$

$$\kappa_{m2} = v_{m20} + v_{m2S}\ S_0 \quad \text{(switch of BT to slow growth mode, boosted by mucin)} \tag{12}$$

##### Nutrient concentrations

The concentrations of nutrients change over time due to their consumption for bacterial growth (rates $\gamma$) and to their production by the bacteria (rates $\alpha$):

$$\text{Glucose:} \quad \frac{dS_1}{dt} = -\gamma_{11}\ \mu_{11}\ X_1 - \gamma_{21}\ \mu_{21}\ X_2 \tag{13}$$

$$\text{Acetate/lactate:} \quad \frac{dS_2}{dt} = \alpha_{12}\ \mu_{11} X_1 + \alpha_{22}\ \mu_{21}\ X_2 - \gamma_{12m}\ \mu_{m12}\ \hat{X}_1 \tag{14}$$

$$\text{Mucin sugars:} \quad \frac{dS_3}{dt} = \alpha_{13}\ S_0\ \bar{X}_1 + \alpha_{23}\ S_0\ \bar{X}_2 - \gamma_{13}\ \mu_{13}\ X_1 - \gamma_{23}\ \mu_{23}\ X_2 \tag{15}$$

$$\text{Butyrate:} \quad \frac{dS_4}{dt} = \alpha_{14}\ \mu_1\ X_1 + \alpha_{14m}\ \mu_{m1}\ \hat{X}_1 \tag{16}$$

#### pH (phenomenologic functions)

pH is not explicitely modeled. To account for the negative effect of pH (assumed to depend exclusively on acetate/lactate) on the growth, we use the following phenomenologic Hill-like functions:

$$\phi_1 = \frac{K_1^m}{K_1^m + S_2^m} \tag{17}$$

$$\phi_2 = \frac{K_2^m}{K_2^m + S_2^m} \tag{18}$$

#### Parameters

Parameter values have been adjusted manually in order to qualitatively reproduce the observed dynamics (Table S2).

#### Observations

The time series generated by numerical simulations of the model are shown in Figures S1 (in absence of mucin) and S2 (in presence of mucin).

In absence of mucin:

- The two peaks of BT in monoculture can be reproduced: the first peak is due to growth on glucose. The second peak results from growth on WC residual sugars, starting when glucose is sufficiently low. After depletion of glucose and other sugars, BT goes to zero.
- The long-standing survival of RI in monoculture is achieved through maintenance in the slow growth state. After depletion of glucose and of WC residual sugars, the amount of growing bacteria goes to zero but a certain fraction converges to the slow growth state. In this slow growing state, RI consumes acetate/lactate.
- In the RI/BT co-culture, a larger growth peak is reached by BT (which better exploits sugars resources, but then BT converges to zero, whereas a small fraction of RI switches to the slow growth mode and survives for a longer period of time.
- pH negatively affects BT growth more than RI growth.

In presence of mucin:

- The long-term survival of BT in monoculture is achieved by the continuous growth on mucin sugars and by the mucin-boosted switch to slow growth mode. This hypothesis was introduced to explain the long term survival of BT observed only in presence of mucin.
- RI first grows on glucose and then, when glucose is depleted, on mucin sugars. It is however not sufficient to compensate death. Because acetate/lactate and mucin sugars are always present, the switch to the slow growth mode is not induced.
- The same dynamics happens in RI/BT co-culture: BT survives for a long period of time due to the slow growth state (boosted by mucin), whereas RI is out-competed.
- pH dynamics/inhibition remains unchanged compared to the case without mucin.

| Name | Matlab | Description | Value |
|---|---|---|---|
| $S_0$ | S0 | mucin (absent or present) | 0 or 1 |
| $\delta_1$ | d1 | death rate of RI | 0.2 |
| $\delta_2$ | d2 | death rate of BT | 0.2 |
| $f_{X1}$ | f1 | fraction of attached RI (*) | 0.25 |
| $f_{X2}$ | f2 | fraction of attached BT (*) | 0.25 |
| | | (*) may also include a small fraction of liquid bacteria that degrade non-monomeric mucin products spontaneously released from the beads | |
| $v_{11}$ | v11 | growth rate of RI on glucose | 0.6 |
| $K_{11}$ | K11 | Monod constant for growth of RI on glucose | 0.5 |
| $v_{13}$ | v13 | growth rate of RI on mucin sugar | 0.1 |
| $K_{13}$ | K13 | Monod constant for growth of RI on mucin sugar | 0.5 |
| $v_{12m}$ | v12 | growth rate of RI on acetate (slow growth mode) | 0.01 |
| $K_{12m}$ | K12 | Monod constant for growth of RI on acetate/lactate | 0.1 |
| $v_{21}$ | v21 | growth rate of BT on glucose | 0.7 |
| $K_{21}$ | K21 | Monod constant for growth of BT on glucose | 0.5 |
| $v_{23}$ | v23 | growth rate of BT on other mucin sugar | 0.6 |
| $K_{23}$ | K23 | Monod constant for growth of BT on other mucin sugar | 0.05 |
| $K_{i11}$ | Ki11 | inhibition constant of RI growth on glucose by mucin sugar | 1 |
| $K_{i13}$ | Ki13 | inhibition constant of BT growth on mucin sugars by glucose | 0.1 |
| $v_{m1}$ | vm1 | rate of switch from growing RI to slow growth mode | 0.04 |
| $k_{i1}$ | Ki1 | rate of switch from growing RI to slow growth mode | 0.1 |
| $k_{i3}$ | Ki3 | rate of switch from growing RI to slow growth mode | 0.6 |
| $h$ | h | Hill coefficient for switch of RI to slow growth mode | 4 |
| $v_{m20}$ | vm20 | basal rate of switch to slow growth mode | 0.001 |
| $v_{m2S}$ | vm2S | rate of switch to slow growth model for BT boosted by mucin | 0.1 |
| $\delta_{m1}$ | dm1 | death rate of RI in slow growth state | 0.004 |
| $\delta_{m2}$ | dm2 | death rate of BT in slow growth state | 0.005 |
| $\gamma_{11}$ | g11 | consumption rate (1/yield) of glucose by RI | 2 |
| $\gamma_{13}$ | g13 | consumption rate (1/yield) of mucin sugar by RI | 2 |
| $\gamma_{12m}$ | g12m | consumption rate (1/yield) of acetate by RI (slow growth) | 2 |
| $\gamma_{21}$ | g21 | consumption rate (1/yield) of glucose by BT | 2 |
| $\gamma_{23}$ | g23 | consumption rate (1/yield) of mucin sugar by BT | 0.5 |
| $\alpha_{12}$ | a12 | production rate of acetate by RI | 1 |
| $\alpha_{22}$ | a22 | production rate of acetate by BT | 1 |
| $\alpha_{13}$ | a13 | production rate of mucin sugar by RI | 2 |
| $\alpha_{23}$ | a23 | production rate of mucin sugar by BT | 2 |
| $\alpha_{14}$ | a14 | production rate of butyrate by RI | 2 |
| $\alpha_{14m}$ | a14m | production rate of butyrate by RI (slow growth mode) | 2 |
| $K_1$ | Ki1 | threshold of sensitivity of RI growth to pH | 0.8 |
| $K_2$ | Ki2 | threshold of sensitivity of BT growth to pH | 0.6 |
| $m$ | m | Hill coefficient used in pH functions | 2 |

Table S2: Parameters.

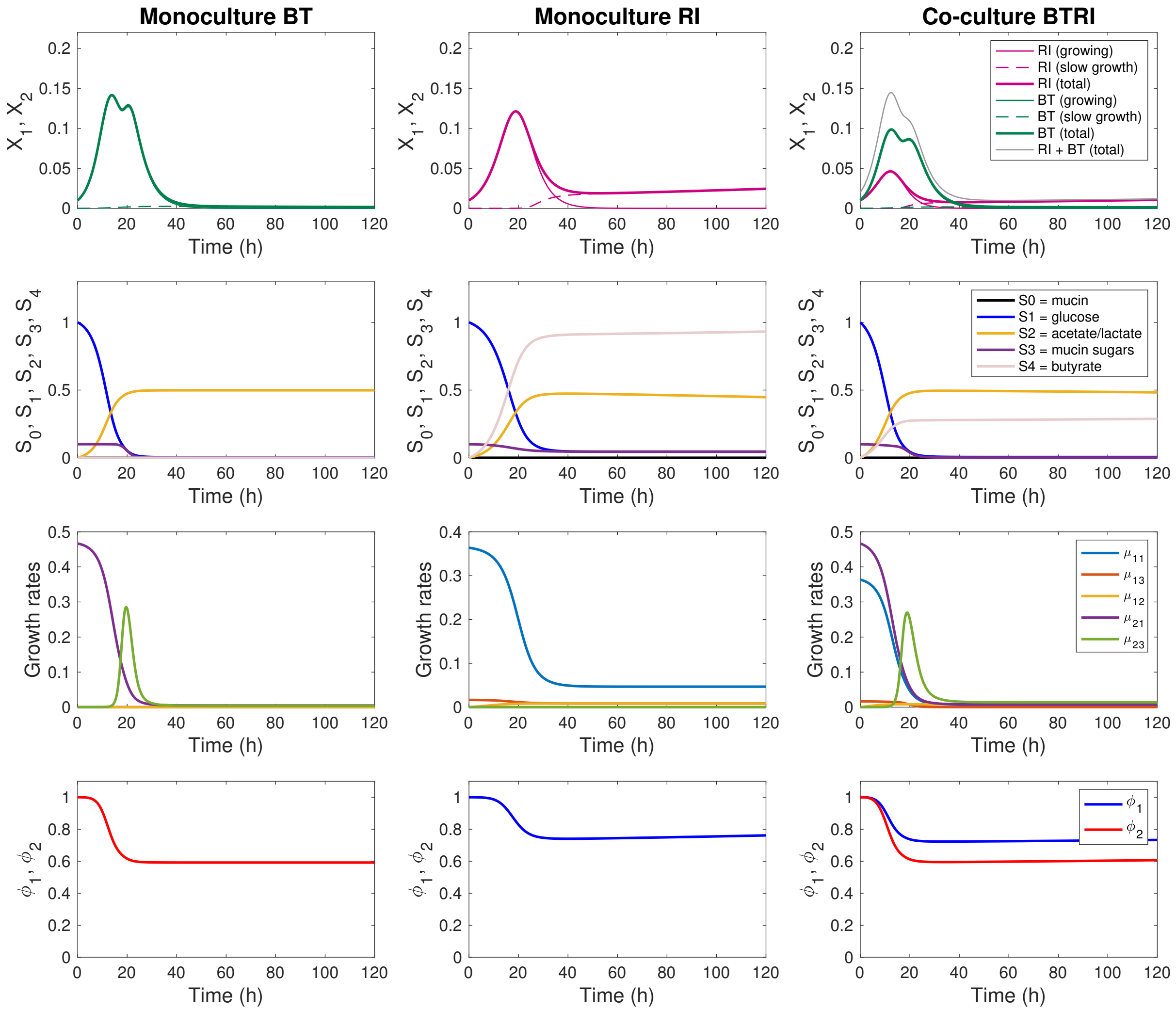


Figure S1: Growth curves in absence of mucin beads ($S_0 = 0$).

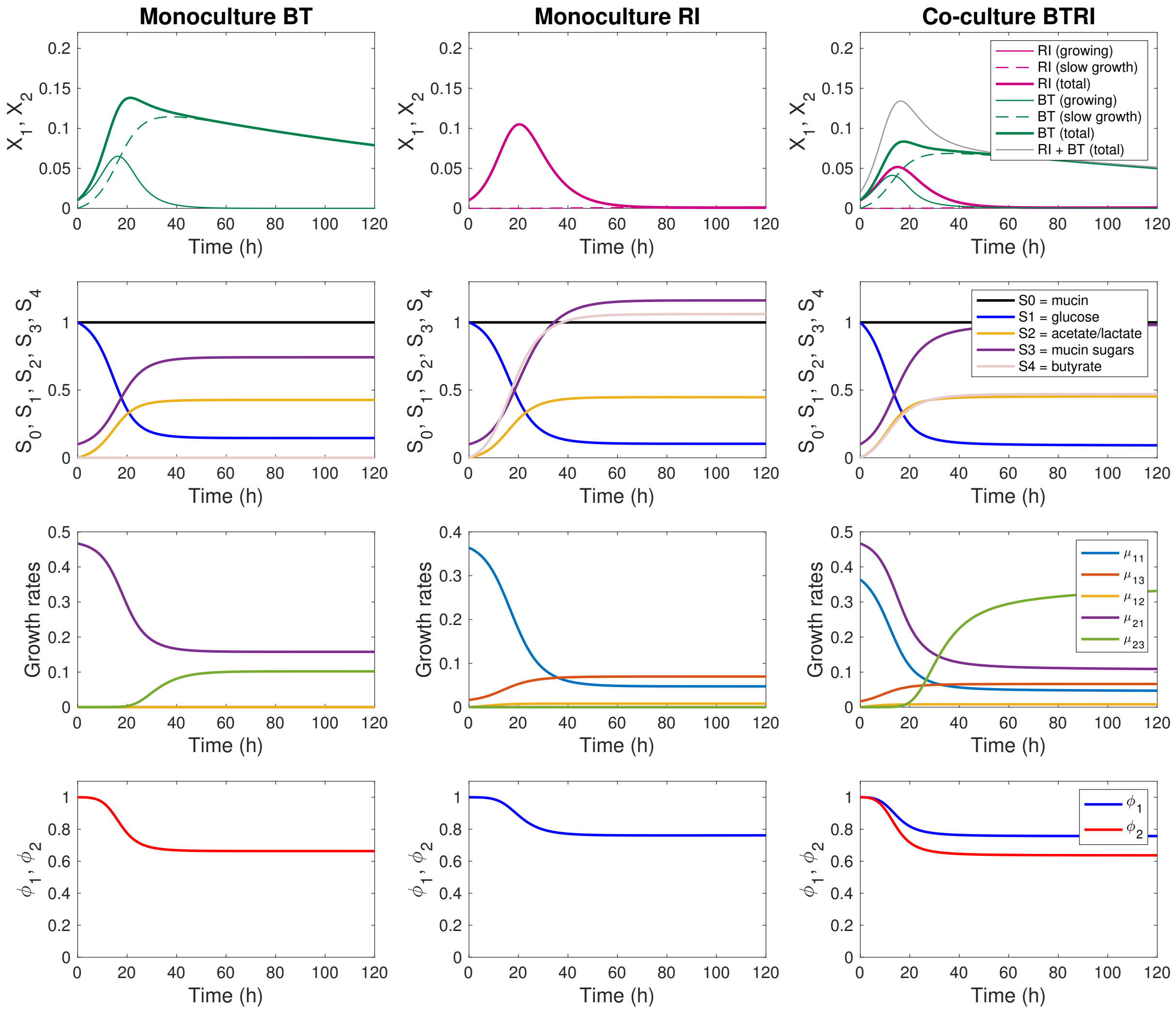


Figure S2: Growth curves in presence of mucin beads ($S_0 = 1$).
